# Cofea: correlation-based feature selection for single-cell chromatin accessibility data

**DOI:** 10.1101/2023.06.18.545397

**Authors:** Keyi Li, Xiaoyang Chen, Shuang Song, Lin Hou, Shengquan Chen, Rui Jiang

## Abstract

Single-cell sequencing technologies have revolutionized the understanding of cellular heterogeneity at an unprecedented resolution. However, the high-noise and high-dimensional nature of single-cell data poses challenges for downstream analysis, and thus increases the demand for selecting biologically informative features when processing and analyzing single-cell data. Such approaches are mature for single-cell RNA sequencing (scRNA-seq) data, while for single-cell chromatin accessibility sequencing data, the epigenomic profiles at the cellular level, there is a significant gap in the availability of effective methods. Here we present Cofea, a correlation-based framework that focuses on the correlation between accessible chromatin regions, to accurately select scCAS data’s features which are highly relevant to biological processes. With various simulated datasets, we quantitively demonstrate the advantages of Cofea for capturing cellular heterogeneity of imbalanced cell populations or differentiation trajectories. We further demonstrate that Cofea outperforms existing feature selection methods in facilitating downstream analysis, particularly in cell clustering, on a wide range of real scCAS datasets. Applying this method to identification of cell type-specific peaks and candidate enhancers, pathway enrichment analysis and partitioned heritability analysis, we show the potential of Cofea to uncover functional biological process and the genetic basis of cellular characteristics.

## INTRODUCTION

Single-cell sequencing technologies have revolutionized the understanding of cellular heterogeneity (1), heritable phenotypes (2), cell fate determination (3) and many others (4,5) at an unprecedented resolution. The majority of single-cell data is derived from single-cell RNA sequencing (scRNA-seq) and single-cell chromatin accessibility sequencing (scCAS) technologies, which study transcriptomic and epigenomic profiles of individual cells, respectively. In a typical single-cell experiment, 10,000∼1,000,000 features (genes for scRNA-seq data, peaks or bins for scCAS data) will be detected (5–12), while the majority of features are of low signal-to-noise ratio or not relevant to underlying biological processes. The utilization of the whole set of features is destined to impede accuracy and computational efficiency of downstream analysis, and consequently, selecting informative features becomes a crucial step for preprocessing single-cell data.

The development of feature selection methods has been mature for scRNA-seq data. These methods can be divided into three major categories: (i) approaches based on genetic variance across all cells (12); (ii) approaches based on Gini index of gene expression (13,14); and (iii) approaches based on the relationship between dropout events and gene expression (15,16). By contrast, methods dedicated to scCAS data are still in their infancy, and the close-to-binary nature of scCAS data impede the direct applicability of methods for scRNA-seq data to scCAS data. Additionally, scCAS data have other assay-specific characteristics, especially extreme sparsity and tens of times higher dimensions than scRNA-seq data, suggesting the pressing demand for feature selection methods specifically designed for scCAS data.

The mainstream operation for scCAS data analysis is to select features that have at least one read count in cells exceeding a specified number (2,17-20). Signac (21) and epiScanpy (22), the widely-used scCAS data analysis tools in R and Python respectively, also include feature selection approaches in their pipelines. Specifically, Signac selects features with higher degree of accessibility after normalization, while epiScanpy aims to retain chromatin regions that are accessible in approximately half of the cells. In essence, the above methods only focus on the level of accessibility from the perspective of individual features, but disregard the interrelationships among them. Features selected by these methods tend to prioritize information derived from cell types with a substantial number of cells, and consequently, may fail to recognize the rare cell type. In summary, there is still a significant gap in effective feature selection that captures cellular heterogeneity of cells in a scCAS dataset.

To fill this gap, we developed a correlation-based framework, Cofea, to select biologically informative features of scCAS data via placing emphasis on the correlation among features. Unlike existing methods that simply aggregate values of a peak-by-cell matrix to obtain the significance of features, Cofea (i) obtains a peak-by-peak correlation matrix after a stepwise preprocessing approach, (ii) establishes a fitting relationship between the mean and mean square values of correlation coefficients to reveal a prevailing pattern observed across the majority of features, and (iii) selects features that deviate from the established pattern to facilitate a more nuanced preservation of the underlying biological complexities. Through various simulated datasets, we quantitatively demonstrated advantages of Cofea for selecting informative features that highly relate to not only balanced or imbalanced cell groups, but also continuous differentiation trajectories. When applied to 54 real scCAS datasets with different tissues, dimensions and qualities, Cofea showed notable and stable performance to facilitate dimensionality reduction and cell clustering, the key tasks of downstream analysis. Features selected by Cofea contain more cell type-specific peaks and candidate enhancers than baseline methods, suggesting the superior relevance to underlying biological processes. Utilizing these features to pathway enrichment analysis and partitioned heritability analysis demonstrates the potential of Cofea in uncovering functional biological process and provides insights into the genetic basis of cellular characteristics. In addition, systematic ablation and dropout experiments further illustrates the robustness and reliability of Cofea.

## MATERIAL AND METHODS

### The framework of Cofea

Cofea is a correlation-based method designed to identify a subset of features that exhibit significant heterogeneity across cells, thereby facilitating downstream analysis of scCAS data. In scCAS data analysis, the data is commonly represented as a peak-by-cell matrix, with each peak treated as a feature. Given a raw peak-by-cell scCAS count matrix ***X*** ∈ ℝ^*p*×*n*^ (*p* features and *n* cells) as input, Cofea outputs significance scores of all peaks and ranks them accordingly. If the user also inputs the number of selected features, Cofea will additionally output a processed matrix that includes only the selected features based on significance scores. As shown in Figure 1, Cofea contains two major steps for processing scCAS data. First, Cofea applies a series of stepwise preprocessing operations to the peak-by-cell matrix, including term frequency-inverse document frequency (TF-IDF) transformation and cell-wise principal component analysis (PCA) transformation. These operations aim to normalize the peaks and mitigate the impact of imbalanced cell populations within a scCAS dataset. Second, Cofea employs a correlation-based evaluation workflow to assess peaks based on the pattern of correlation coefficients. More specifically, the workflow focuses on the mean and mean square values of inter-peak correlation coefficients (i.e., correlation coefficients between different peaks), and utilizes a locally weighted scatterplot smoothing (LOWESS) model to capture the inherent pattern of the relationship between these values. The degree of deviation from this pattern then serves as the importance score of a peak, with outliers being defined as "informative peaks". By leveraging the dimensionality reduction approach at the cell level and a parallel computing strategy, Cofea effectively reduces computational complexity and memory usage, enhancing the reliability and efficiency of utilizing computing resources.

**Figure 1.**
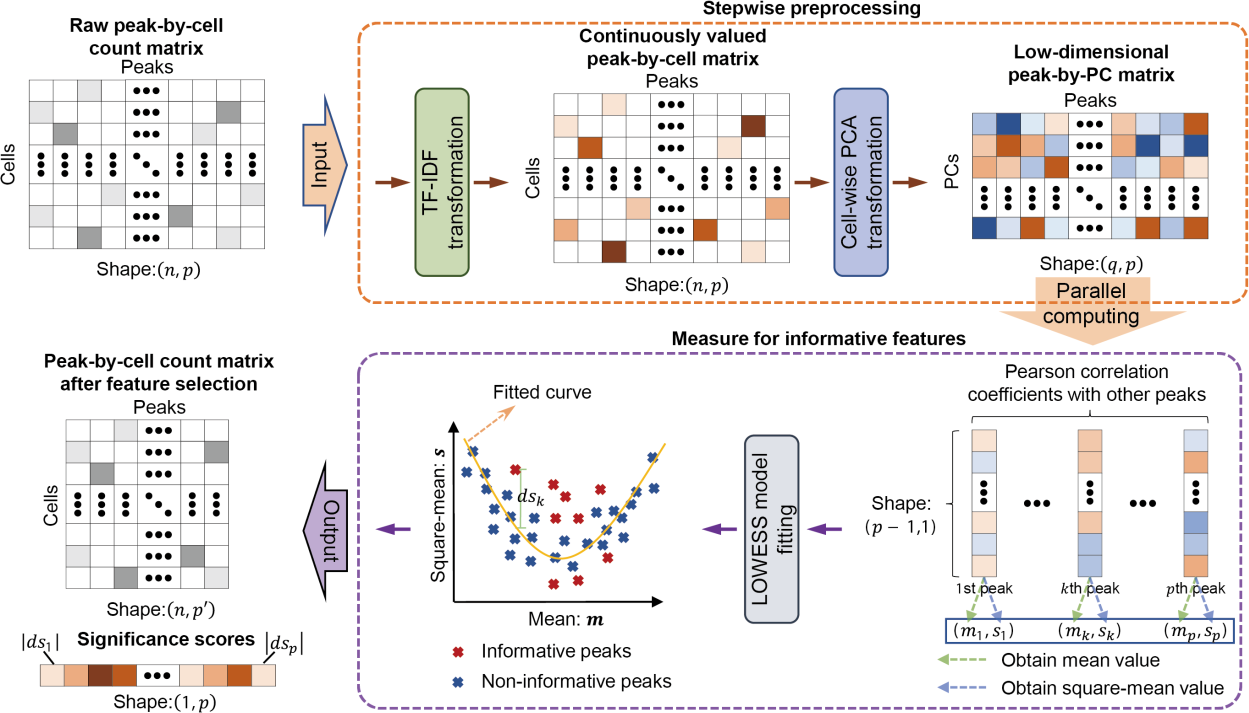
Overview of Cofea. Cofea’s feature selection includes two main steps, namely Stepwise preprocessing and Measure for informative features. First, Cofea receives input in the form of peak-by-cell matrix, and performs TF-IDF transformation and cell-wise PCA transformation to obtain the low-dimensional peak-by-PC matrix. Then, Pearson correlation coefficients between features are calculated, and LOWESS fitting is carried out to get the significance scores for each feature. The features that deviated furthest from the fitting curve are selected, and the index and significance scores of the features are output.

### Stepwise preprocessing in Cofea

Cofea first applies TF-IDF transformation to the input peak-by-cell matrix following Signac (21), a widely used toolkit for scCAS data analysis (23–26). For an input scCAS dataset, considering the element ***x***_***ij***_, which represents the count value of the ***i***th peak of the ***j***th cell in the peak-by-cell matrix ***X***, the TF-IDF transformation could be formulated as follows:

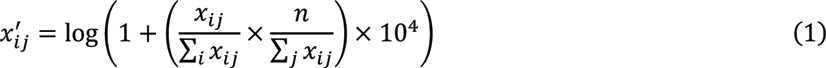

where 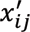 denotes the element associated with *x*_*ij*_ in the newly-formed matrix ***X***′ ∈ ℝ^*p*×*n*^. The TF-IDF transformation converts the read count matrix into a continuously valued matrix by upweighting peaks that do not occur very frequently and down-weighting prevalent peaks, which has shown notable performance in recent studies for scCAS data analysis (2,17,20,21,27). Cofea also provides other two implementations of TF-IDF transformation, which are used in a recent benchmark (referred to as TF-IDF_origin) and scOpen (referred to as TF-IDF_scOpen) respectively, as alternatives (Supplementary Note S1).

To reduce the computing resources, Cofea also applies cell-wise PCA transformation to the matrix ***X***′, and then obtains a peak-by-PC matrix ***P*** ∈ ℝ^*p*×*q*^, where *q* is the number of principal components (PCs) and set to 100 as the default value. This operation not only ensures the adaptability and robustness of Cofea with various numbers of cells, but also improves the computational efficiency.

### Evaluation workflow for informative features

The evaluation for informative features in Cofea relies on a peak-by-peak correlation matrix ***C*** ∈ ℝ^*p*×*p*^, which could be obtained by computing Pearson correlation coefficients (PCC) between peaks as follows:

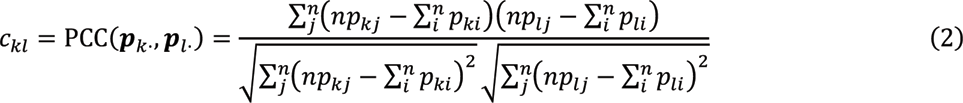

where *p*_*kj*_ represents the element in ***P***, and *c*_*kl*_ is an element in ***C***, representing the correlation between the *k*th peak (denoted as the row vector ***p***_*l*·_) and the *l*th peak (denoted as the row vector ***p***_*l*·_) in the PCA-transformed matrix ***P***. Next, a mean vector ***m*** and a mean square vector ***s*** of inter-peak correlation coefficients could be calculated based on column vectors of ***C***. To accomplish this, for each column vector, such as the *j*th vector ***c***_*j*_, Cofea first removes the self-correlation coefficient, of which the index in ***c***_*j*_ is *j* and the value is fixed to 1, and then calculates the mean value and mean square value of the remaining elements to obtain *m*_*j*_ and *s*_*j*_, that is, the element in ***m*** and ***s***, respectively. Note that since the intermediate storage for the whole matrix ***C*** may cause the memory crash due to the large size of *p* × *p*, Cofea only uses a temporary vector to store the column vector ***c***_*j*_ for generating *m*_*j*_ and *s*_*j*_, and then replaces the value in temporary vector with next column vector ***c***_*j*+1_. To accelerate the procedure, Cofea uses a tailored parallel computing strategy for processing multiple column vectors simultaneously (Supplementary Note S2). Furthermore, Spearman correlation coefficients (SPCC) and Cosine similarity coefficients (CSC) are also provided as alternative hyperparameters to obtain correlation coefficients between peaks (Supplementary Note S3).

To learn the relationship between the mean and mean square values of correlation coefficients, Cofea fits a LOWESS model using the paired data in ***m*** and ***s***, such as (*m*_*j*_, *s*_*j*_), and calculates the corresponding residual between *s*_*j*_ and the predicted value 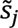 as *ds*_*j*_. The LOWESS model aims to capture the prevailing pattern inherent to the majority of features, and the absolute value of residual |*ds*_*j*_| represents a significance score for the corresponding feature. We use a normal distribution to estimate residuals of all features, and calculate the quantile corresponding to each feature based on the fitted distribution. To mitigate the impact of overfittings due to the outliers, Cofea carries out model fitting three times, and during each time, filters out data points with the quantiles below 5% and above 95% for next time of model fitting. Finally, Cofea ranks the features via significance scores derived from the last fitted LOWESS model, and identifies a user-defined number of features as “informative features” (also denoted as “informative peaks”).

### Data collection and preprocessing

#### Simulated datasets

The performance of feature selection for scRNA-seq data can be assessed quantitatively by comparing with marker genes (28,29). However, the quantitative evaluation of feature selection methods for scCAS data poses a challenge due to the absence of well-defined annotations for marker features of scCAS data. To this end, we generated five high-fidelity scCAS datasets (Dataset S1 to S5, details in Supplementary Note S4) with ground truths of informative peaks, enabling a quantitative evaluation of feature selection methods. All the simulated datasets preserve characteristics as with real scCAS data, including high-sparsity, high-dimensional and close-to-binary nature.

For each dataset, the peak-by-cell matrix contains two types of peaks: cell type-specific peaks and background peaks. The cell type-specific peaks are characterized by higher accessibility signals in the corresponding cell type than others, while the background peaks have the consistent level of accessibility across different cell types. We then used Cofea to identify peaks with the same number of all cell type-specific peaks, and calculated the overlapped proportion between cell type-specific peaks and the selected peaks as a metric. For each cell type, a higher overlap value indicates a superior identification performance in capturing the cellular heterogeneity of the corresponding cell type.

#### Real datasets collection and preprocessing

To evaluate the performance of Cofea, we collected four previously published datasets including Buenrostro2018 (5), Joshua2021 (6), Neurips2021 (7), Sai2020 (8) datasets, and three atlases, including mouse cell atlas (MCA) (10), fetal cell atlas (FCA) (9) and human genome cell atlas (HGCA) (11). Each atlas consists of multiple datasets, with each individual dataset representing a specific tissue or organ. We denote each dataset within the atlas by combining the atlas name with the name of the corresponding tissue or organ. For instance, the MCA heart dataset specifically represents the heart tissue within the MCA atlas. Following the recommendation by several studies of scCAS data analysis (2,17-20), we conducted data preprocessing by filtering out the peaks that have at least one read count in less than 1% of cells. A summary of the above datasets is shown in Supplementary Table S1.

### Model Evaluation

Feature selection is a crucial step for many downstream analysis tasks in scCAS data analysis (19,21,22), while systematically benchmarking the performance of related methods remains a tough challenge. This is primarily due to the limited knowledge about the congruent relationship between cell types and accessible chromatin regions, rendering it impracticable to directly calculate the accuracy of identified features. With this consideration, we conducted a comprehensive evaluation to illustrate the benefits of Cofea from the following two perspectives: (i) whether features provided by Cofea can outperform existing feature selection methods in facilitating downstream analysis; (ii) whether informative features themselves identified by Cofea can reveal biological insights.

From the perspective of facilitating downstream analysis, we conducted dimensionality reduction and cell clustering, the two pervasive downstream analytic tasks which can be quantitatively evaluated by targeted assessment metrics, to measure the performance of feature selection. We utilized Signac (21), one of the most commonly used tools for scCAS data analysis, to perform dimensionality reduction and cell clustering. During the Louvain clustering in Signac, we adopt a binary search approach for identifying an appropriate resolution, to make the number of clusters closely match the number of cell types. By using an equal number of features for comparison, all the metrics intuitively and quantitatively assessed the impact of feature selection on downstream analytic tasks, offering insights into the accuracy and quality of the informative features identified by different methods.

From the perspective of revealing biological insights, we evaluated Cofea via several numeric experiments including cell type-specific peaks annotation, candidate enhancers identification, functional pathway enrichment and partitioned heritability analysis.

First, we quantified the performance of Cofea by measuring the overlapped proportion of the set of identified informative peaks and cell type-specific peaks predicted by the tl.rank_features function in epiScanpy. To be more specific, if cell labels are provided, the function will outputs cell type-specific peaks that exhibit differential accessibility across different cell types. For each cell type, a higher proportion of overlap indicates a stronger association between the identified features and the corresponding cell type.

Second, we collected candidate enhancers from the scEnhancer database (30). We employed the liftOver (31) to convert the coordinates of the identified peaks from GRCm38 (UCSC mm10) to GRCm37 (UCSC mm9), ensuring the compatibility with enhancers. Similarly, we calculated the overlapped proportion of enhancers and features identified by Cofea. A higher proportion of overlap indicates a greater relevance between the identified peaks and regulatory patterns to the corresponding cell type.

Third, we focused on the set of features uniquely identified by Cofea. We clustered these features using Louvain clustering algorithm based on the peak-by-PC matrix generated by the intermediate step of Cofea. To gain functional insights into these features, we performed pathway enrichment on each cluster of features using the Genomic Regions Enrichment of Annotations Tool (GREAT) (32), and compared the enriched pathways with cellular functions of cells in a scCAS dataset, to detect a deeper understanding of the biological functions encoded within these features.

Fourth, we analysed the heritability of five brain-related phenotypes within specific partial peaks using partitioned linkage disequilibrium score regression (S-LDSC) (33,34). To achieve this objective, we first mapped the genome coordinates of the peaks identified by Cofea to GRCh37/hg19. S-LDSC was then applied to each phenotype to estimate the enrichment of heritability within the identified peaks. To perform the analysis, we followed the recommended LDSC workflow, using HapMap3 SNPs and European samples from the 1000 Genomes Project as the LD reference panel. The GWAS summary statistics were downloaded from https://data.broadinstitute.org/alkesgroup/sumstats_formatted/.

### Metrics for assessment of dimensionality reduction and cell clustering

We adopted four widely-used metrics for assessing the clustering results as recommended in previous studies (35–37), namely normalized mutual information (NMI), adjusted mutual information (AMI), adjusted Rand index (ARI), and homogeneity score (Homo). A higher value approaching 1 for these metrics indicates a better performance of cell clustering, indicating the effectiveness of the feature selection method. Additionally, we utilized another two metrics suggested in scIB (35) for evaluating the conservation of biological variance: the average silhouette width (ASW) and cell-type LISI (cLISI) (38). ASW assesses the density and separation of clusters and cLISI measures the accuracy of an embedding with cell-type prediction. Similarly, higher values of ASW and cLISI are indicative of better feature selection performance. The detailed descriptions and formulas of the above metrics are available in Supplementary Note S5.

### Baseline methods

We compared the performance of Cofea with three baseline methods: selecting features with the highest degree of accessibility (referred to as HDA), and two scCAS data analysis toolkits, epiScanpy (22) and Signac (21). HDA is the most widely used feature selection method for scCAS data analysis (2,17-19), as it selects peaks that have at least one read count in most cells. We implemented HDA in assessment Python following its instruction. Meanwhile, epiScanpy and Signac both include feature selection as a fundamental step in their pipelines. Source code for implementing the two methods was obtained from their studies. The detailed descriptions of the above three methods are available in Supplementary Note S6.

It is worth noting that we also conducted a series of experiments to evaluate the effectiveness of scRNA-seq methods, including highly variable genes (HVG) (12), M3drop, NBdrop (15), and GiniClust (13), in scCAS data analysis (Supplementary Note S7 and Supplementary Figure S1). Our results demonstrated that these scRNA-seq methods are not suitable for effectively scaling to scCAS data analysis and thus we did not include them as baseline methods in our comparison.

## Results

### Cofea accurately identifies cell type-specific peaks in simulated datasets

Peaks, the chromatin regions with high level of accessibility, are commonly used features in scCAS data analysis (39,40). However, evaluating feature selection methods for scCAS data poses significant challenges due to the lack of ground truths of informative peaks. Here we generated five simulated scCAS datasets as with the characteristics of real-world data, and manually set cell type-specific peaks as the ground truths of informative peaks (MATERIALS AND METHODS). To provide a clear display of cell populations and accessibility level of peaks in simulated datasets, we visualized the peak-by-cell matrices of simulated datasets with part of peaks and cells, and found that the simulated data align well with the intended design outlined in the Methods section (Figure 2A). By utilizing the uniform manifold approximation and projection (UMAP), we generated a low-dimensional representation of cells to show the inherent cellular manifold. The simulated cells within datasets S1-S4 exhibit distinct and discrete cell populations, while dataset S5 is characterized by cells from a continuous differentiation trajectory.

**Figure 2.**
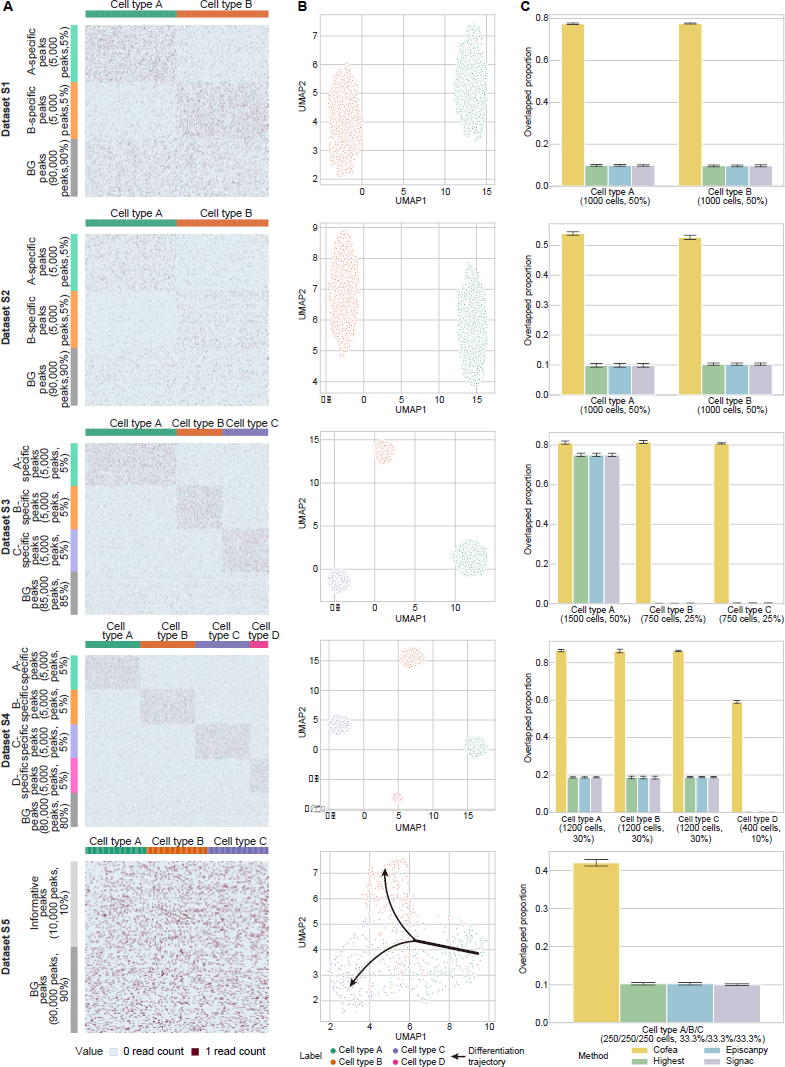
Evaluation of cell type-specific peak identification performance using five simulated datasets. A, Visualization of simulated datasets. Note that for each dataset, we only showed 200 type-specific peaks for each cell type and 200 background peaks, by random sampling without replacement, and then randomly selected 20% cells for each cell type for visualization without replacement. In each subplot to the corresponding dataset, each row denotes a peak and each column denotes a cell. Here BG means background, and A-specific represents peaks that are specific in cell type A. B, UMAP visualization of five simulated datasets, using the low-dimensional representation of cells via the pipeline in Signac. C, Overlapped proportion of the informative peaks identified by Cofea and baseline methods with cell type-specific peaks preset in each simulated dataset. We generated each simulated dataset five times with different random seeds, and then performed feature selection using Cofea and baseline methods. The measure of center for the error bars denotes the average of overlapped proportion for different repeated experiments, and the error bar denotes the estimated standard error in five bootstrap samples. From top to bottom are the results of datasets S1-S5.

We then calculated the overlapped proportion between cell type-specific peaks and identified peaks as a metric for model evaluation, and benchmarked Cofea against three competing feature selection methods widely used in scCAS data, namely HDA, epiScanpy and Signac (MATERIALS AND METHODS). As shown in Figure 2C, Cofea achieved a notable and stable performance for cell type-specific peaks identification, while the performance of baseline methods fluctuated across different cell types in different datasets. On dataset S1, Cofea identifies a higher proportion of informative features that overlap with cell type-specific peaks, compared to baseline methods. This suggests that when coping with features with a consistent degree of chromatin accessibility, Cofea has the potential to detect features with higher cellular heterogeneity. On datasets S2 and S3, of which peaks have a dynamic range of chromatin accessibility, Cofea also outperforms other methods. As the peak-calling process only focuses on accessibility signals of aggregated single cells, accessible chromatin sites related to rare cell types are routinely neglected (41). To test the performance of Cofea in uncovering cellular heterogeneity of imbalanced cell populations, we used dataset S4, which consists of both common cell types (A, B and C) and a rare cell type (D). The results illustrated that Cofea has superior performance, while three competing methods failed to recover rare cell types. We further validated Cofea in dataset S5 with the manifold of tree-structure, and consistently, features identified by our model are more relevant to differentiation trajectories. To summarize, Cofea can not only accurately identify the informative features that highly relevant to balanced or imbalanced cell populations with various signals of accessibility, but can also discover accessible chromatin sites specifically for cell differentiation.

### Cofea facilitates dimensionality reduction and cell clustering of scCAS data

Cell type annotation is an essential step for single-cell data analysis. The typical method is dimensionality reduction and cell clustering, followed by assigning the putative cell type label to each cluster (40). We hereby used the two fundamental tasks for scCAS analysis, to evaluate the performance of Cofea in real scCAS datasets (MATERIALS AND METHODS).

At the outset, we sought to detect the minimal number of features selected by Cofea required to satisfy the accuracy requirements of downstream analysis. We conducted a preliminary experiment on datasets from three cell atlases. Specifically, we varied the number of features selected by Cofea and baseline methods, and performed dimensionality reduction and cell clustering to obtain predicted labels. Given cell labels as ground truth, we evaluated the performance of the methods by NMI, which is preferred to ARI because the sizes of cell populations are imbalanced and when there are rare cell types (37). As shown in Figure 3A, using Cofea as the feature selection method, clustering performance consistently surpasses that using baseline methods among different numbers of features. As the number gradually increases, the performance of cell clustering improves. On the basis of preserving its superiority, Cofea achieves comparable clustering performance with the least feature number to that of using the whole set of features. More specifically, 20,000 features selected by the Cofea proves to be adequate for downstream analysis to maintain the accuracy, while baseline methods fall short in this scenario.

**Figure 3.**
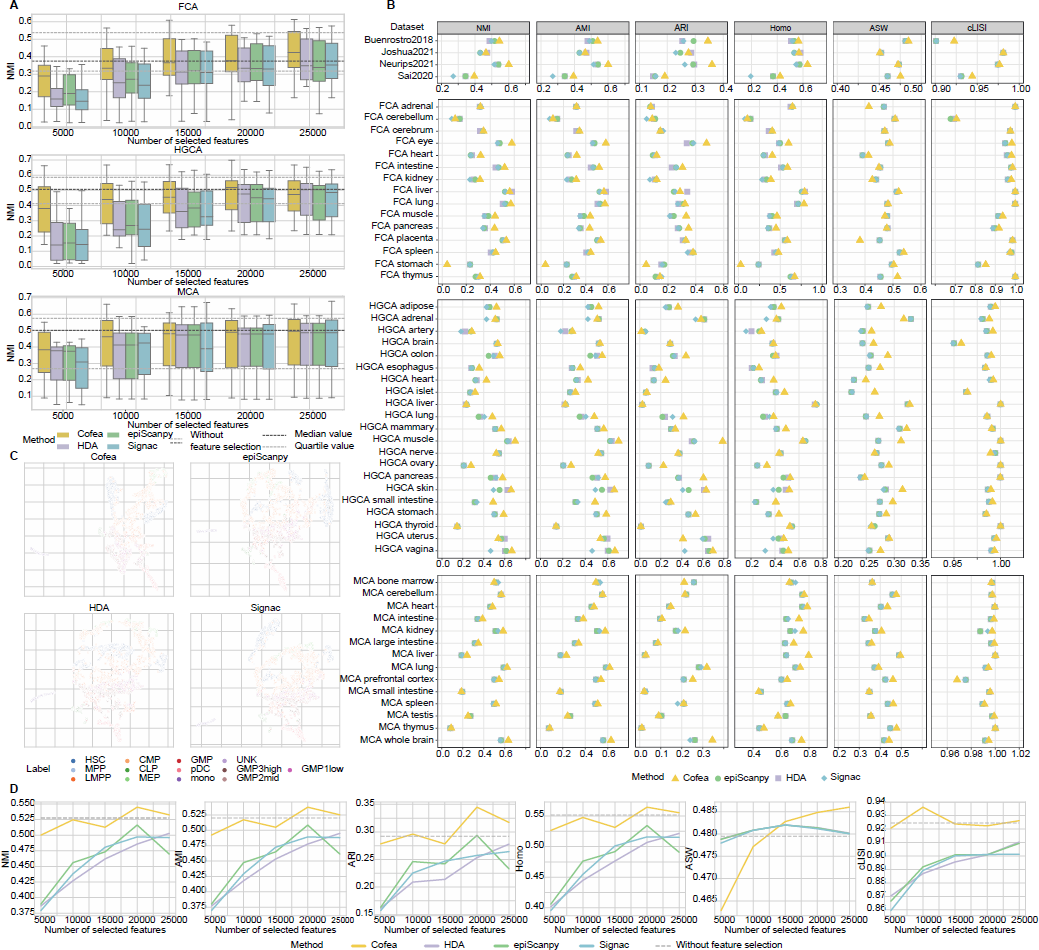
Evaluation of feature selection performance via dimensionality reduction and cell clustering in real scCAS datasets. A, Cell clustering performance evaluated by NMI scores using different number of features selected by Cofea, HDA, epiScanpy and Signac, respectively, on datasets in MCA, FCA and HGCA atlases. For each feature selection method, we set 5000, 10000, 15000, 20000 and 25000 as the number of selected features. The measure of center for the error bars denotes the median value of NMI scores on different datasets in an atlas, and the error bar denotes the maximum and minimum value after removing the outliers, which are defined as the values whose distance from the median is greater than 1.5 times the quartile distance. The black dotted line denotes the median NMI score of cell clustering without feature selection, and the gray dotted lines denote the first and third quartiles. B, Dimensionality reduction and cell clustering performance on all the datasets we collected, using 20,000 features selected by Cofea, HDA, epiScanpy and Signac, respectively. The performance is evaluated by NMI, ARI, AMI, Homo, ASW and cLISI scores. C, UMAP visualization of cells in the Buenrostro2018 dataset, using 20000 features selected by Cofea, HDA, epiScanpy and Signac, respectively. D, Dimensionality reduction and cell clustering performance on the Buenrostro2018 dataset, using features selected by Cofea, HDA, epiScanpy and Signac, respectively. For each feature selection method, we set 5000, 10000, 15000, 20000 and 25000 as the number of selected features, and the performance is evaluated by NMI, ARI, AMI, Homo, ASW and cLISI scores.

We then fixed the number of selected features to 20,000 and conducted a comprehensive benchmark by incorporating a series of quantitative metrics. We additionally included three clustering metrics, namely AMI, ARI and Homo, and As suggested in a recent benchmark study (35), we also evaluated the performance by utilizing ASW and cLISI (MATERIALS AND METHODS). As shown in Figure 3B, Cofea substantially outperforms other methods across all datasets for promoting cell clustering. HDA and epiScanpy obtain similar even consistent results in most datasets, because the majority of peaks have at least one read count in fewer than 50% cells, leading to 20,000 semblable features identified by the two methods. Regarding the performance of cell clustering and taking NMI as an example, we can find that the features identified by Cofea achieve the highest NMI metric values in 41 datasets (in a total of 54 datasets), indicating the superior performance of Cofea. Similar results were observed for the other five metrics considered in this study. Taking the Buenrostro2018 dataset as an example, we projected cells to 2-dimensional space using UMAP. As shown in Figure 3C, compared to baseline methods, biological variations between cell types are better distinguished when using Cofea for feature selection. For instance, MEP cells and CMP cells are well separated with features selected by Cofea, while other methods markedly mixed the two cell types. To summarize, Cofea can effectively preserve cell heterogeneity and facilitate downstream analysis in practical scenarios of scCAS data analysis.

In addition to comparison measured only by NMI or setting a fixed number of selected features, we also conducted a comprehensive evaluation using all six metrics with various numbers of selected features on all collected datasets (Figure 3D and Supplementary Figure S2-S5). The results demonstrated the superiority and reliability of Cofea, which achieves the overall highest value of metrics in most datasets. Interestingly, we observed a trend when selecting features with a small number such as 5,000, Cofea exhibits significant advantages in facilitating cell clustering, indicating the notable effectiveness from another angle.

### Cofea reveals biological signals by identifying informative features

We next focused on informative features themselves identified by Cofea, from a more intuitive perspective, that is, whether features can reveal biological insights. We chose brain, the most complex part of an animal body, to detect brain-related traits from scCAS datasets. By identifying informative features in an unsupervised manner, Cofea can be used to provide functional insights into the cell populations or tissues, and we demonstrated this from the following perspectives.

First, informative features identified by Cofea can characterize cellular heterogeneity of different cell types. We combined the MCA whole brain, MCA cerebellum and MCA prefrontal cortex datasets to obtain a dataset, referred to as the MCA brain dataset, and selected 5,000 informative peaks of this dataset using Cofea and baseline methods. For each cell type, we employed a built-in function in epiScanpy to annotate 1,000 cell type-specific peaks, which can be considered as marker peaks compared to marker genes in scRNA-seq (Methods). We illustrated the overlapped proportion of cell type-specific peaks and informative peaks in Figure 4A and 4B. We can see for virtually all cell types, the highest proportion lies on the informative peaks identified by Cofea, and the overall performance of Cofea surpasses that of baseline methods, indicating that Cofea better captures the heterogeneity of related cell types.

**Figure 4.**
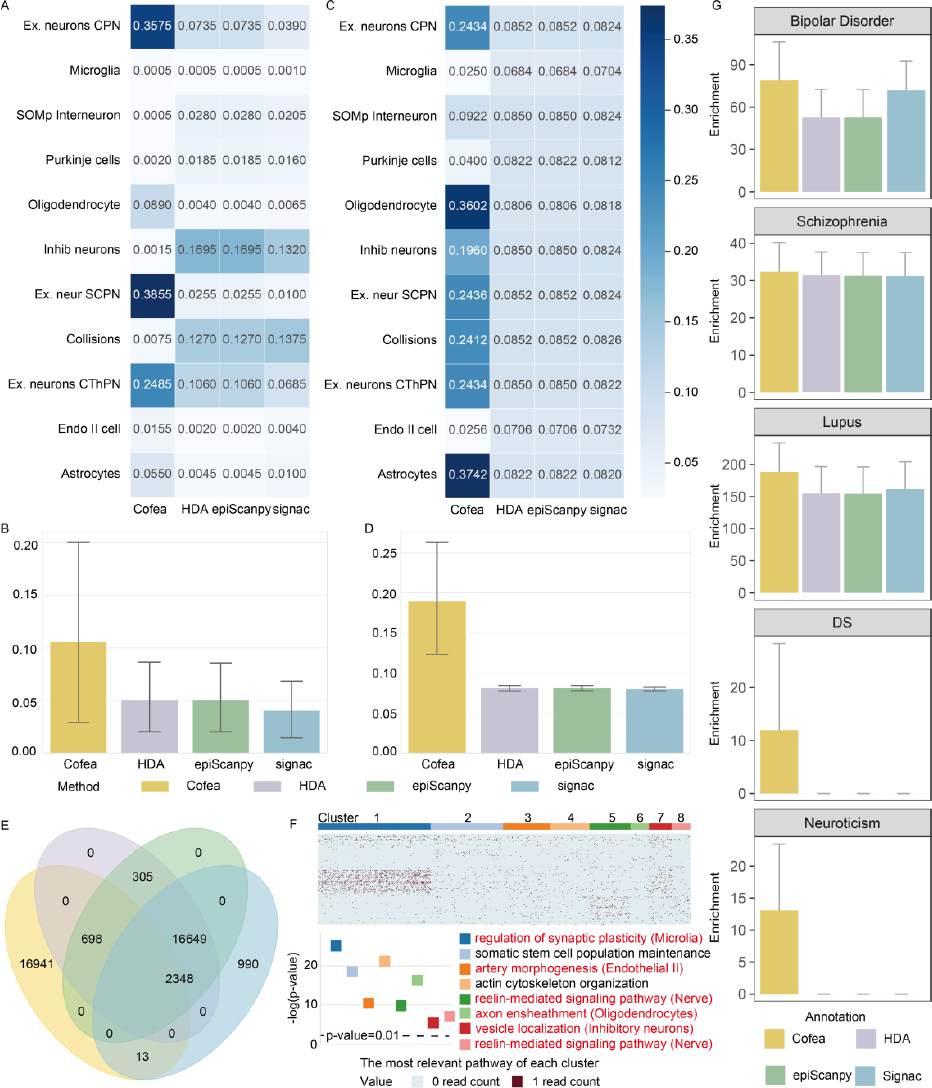
Biological implications of the informative peaks identified by Cofea. A-B, Overlapped proportion of the cell type-specific peaks identified by EpiAnno and the background peaks with cell type specific-peaks related to each cell type (A) or overall cell types (B). C-D, Overlapped proportion of the informative peaks identified by Cofea and baseline methods with brain enhancers collected from scEnhancer related to each cell type (C) or overall cell types (D). In A or C, each row corresponds to a cell type, and each column corresponds to a feature selection method. In B or D, the error bars and centres of error bars represent standard errors and average values of proportion over different cell types. E, Venn diagram of 20,000 peaks selected by Cofea, HDA, epiScanpy and Signac on the MCA Brain dataset. F, Clustering results on the peak-by-PC matrix generated by intermediate procedure of Cofea. Part of cluster of peaks are marked with the most significant pathway enriched by GREAT. The pathway is also associated with a specific cell type corroborated by previous literatures. The bar chart is the Binom Raw p-value corresponding to the biological processes most relevant to each cluster. G, The genetic enrichment of SNPS within the informative peaks identified by Cofea and baseline methods was estimated by stratified LDSC for five brain-related traits. The error bar and the center of the error bar represent the cutter standard error and mean value of enrichment estimation on 200 adjacent SNPs of equal size, respectively.

Second, Cofea can help discover candidate enhancers. scEnhancer provides a single-cell enhancer annotation database and we compared the selected 5,000 informative peaks from the MCA brain dataset with enhancers of brain-related cell types. As the overlapped proportion illustrated in Figure 4C and 4D, peaks provided by Cofea have a high overlapped proportion with enhancers of most cell types, while peaks provided by baseline methods fail to intersect with enhancers. When synthesizing all the cell types, Cofea has significantly immense superiority in tissue-related enhancer detection, suggesting the potential of Cofea to reveal chromatin regulatory landscape in related tissues.

Third, the informative peaks identified by Cofea can help reveal cell type-specific functional implications. We amplified the number of selected features up to 20,000 and identified informative peaks from the MCA brain dataset. An intriguing phenomenon is that the overwhelming majority of peaks (84.705%) identified by Cofea are unique, while the peaks selected by HDA completely overlap with those selected by epiScanpy (Figure 4E). To detect the underlying reasons why the performance of cell clustering with features selected by Cofea can outperform than other methods, we clustered the Cofea-unique peaks to 8 clusters, and performed GREAT analysis (32) to identify the most significant pathway associated with each cluster (MATERIALS AND METHODS, and Figure 4F). It is heartening that, most of the pathways are related to a specific cell type in MCA datasets. To be more specifically, we obtained ‘regulation of synaptic plasticity’ for cluster 1, which is associated with microglia cells (42–44). The enrichment result of cluster 3 is ‘artery morphogenesis’, a relevant function of endothelial II cells (45). Both cluster 5 and cluster 8 are associated with nerve cells (46,47) as peaks of the two clusters are most relevant to ‘reelin-mediated signaling’. Cluster 6 is related to oligodendrocytes (48,49) with the match of the function ‘axon ensheathment’. Cluster 7 corresponds to Inhibitory neurons (50–52), which may be transported by vesicles, and is primarily associated with the biological process of vesicles.

Forth, features selected by Cofea provide insights into the understanding of human traits. We performed partitioned heritability analysis on the features identified by Cofea and other baseline methods with the HGCA brain dataset, on five brain-related phenotypes including Bipolar Disorder, Schizophrenia, Lupus, Down syndrome (DS) and Neuroticism. As shown in Figure 4G, features selected by Cofea have the strongest enrichment of heritability for brain disorders, indicating that Cofea has the potential to preserve genetic information at the tissue level.

### The framework of Cofea is elaborated and has high robustness

As the procedure of Cofea is stepwise and transparent, it provides us an opportunity to reliably examine Cofea in depth. We hereby conducted a series of experiments to demonstrate the stability and robustness of Cofea from two perspectives: (i) the contribution of each module; (ii) the robustness to dropout events and model hyperparameters.

#### Contribution of each module

To validate the contribution of each module, we omitted each part in Cofea in turn but remained the other hyperparameters with default settings, obtaining incomplete variants of Cofea. Taking the Buenrostro2018 dataset as an illustration, we compared cell clustering performance using peaks selected by Cofea and its variants.

First, we obtained a variant by removing the TF-IDF transformation and directly applying the PCA transformation to the raw count matrix. With different number of selected features, Cofea consistently outperforms the variant in all the metrics, suggesting the need of the TF-IDF transformation to accurately characterize scCAS data (Figure 5A and Supplementary Figure S6). Second, we performed a model ablation analysis of omitting the PCA transformation in the stepwise preprocessing of Cofea. By directly using the peak-by-cell matrix to obtain correlation between peaks, the variant achieves comparable clustering performance, while the peak memory use and running time increase by 12.97% and 2404.21%, respectively (Figure 5B and C). Finally, we tested the performance using two other settings in generating PCC between peaks, respectively: (i) we let Cofea regress to generating the whole peak-by-peak correlation matrix, and obtained variant A of Cofea; (ii) we omitted the parallel computing strategy in generating PCC between peaks, and obtained variant B of Cofea. When applying variant A, we encountered memory corruption and failed to select features. As shown in Figure 5D, the computing time of Cofea is 54.42% lower than that of variant B, respectively, indicating that the parallel computing strategy enables Cofea to possess a higher computational efficiency. To summarize, all the above results demonstrated that each module in Cofea is indispensable to achieving effective and efficient feature selection.

**Figure 5.**
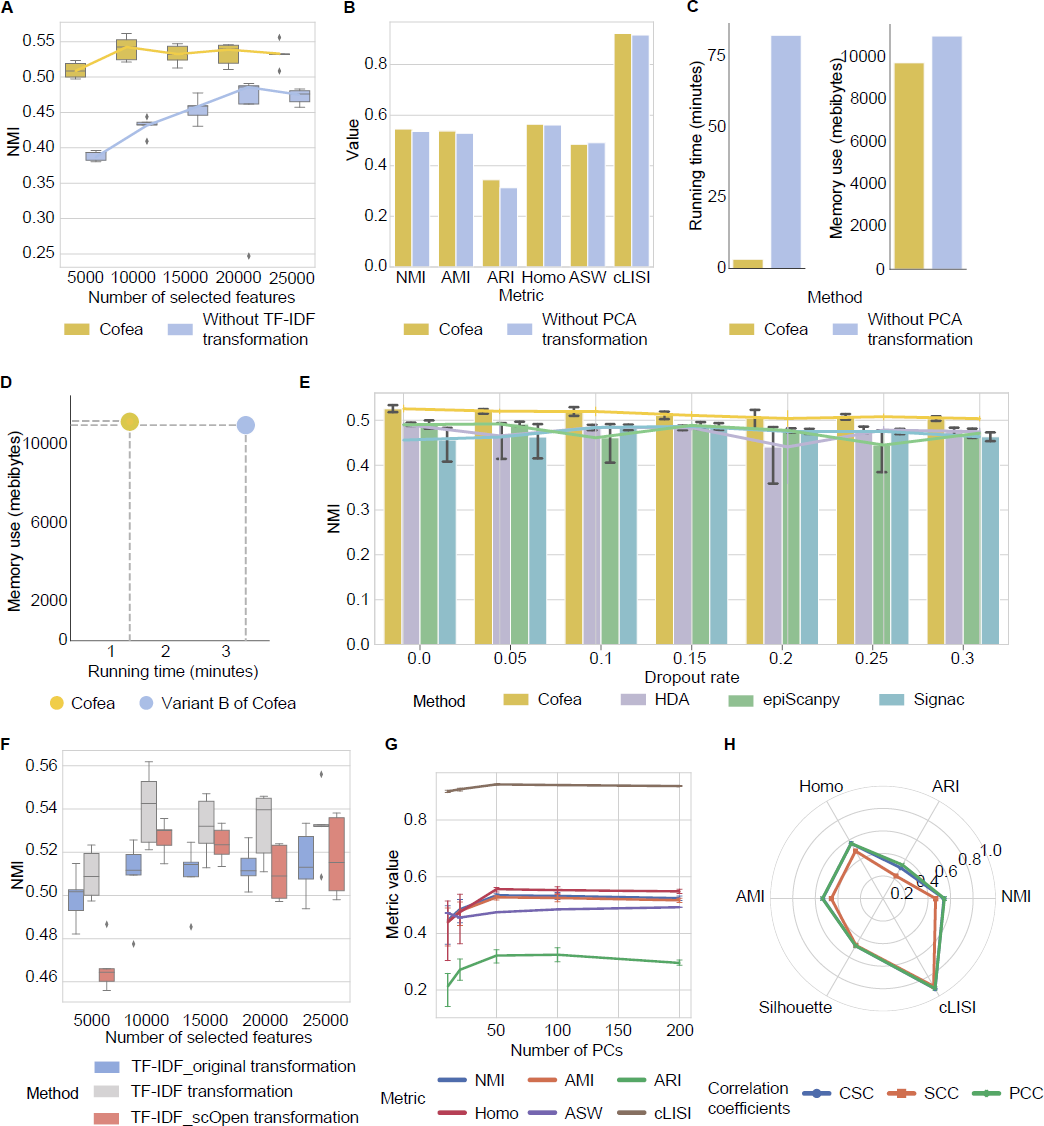
Model ablation and robustness analysis of Cofea. A, Cell clustering performance evaluated by NMI scores with different number of selected features from Cofea and the variant by removing TF-IDF transformation. The measure of center for the boxes and the dot on the line plot are the median value of NMI scores, and the error bar denotes the first and third quartiles. B, Cell clustering performance evaluated by all six metrics (NMI, AMI, ARI, Homo, ASW and cLISI) with 20000 features selected by Cofea and the variant of removing PCA transformation. C, Memory use and running time for feature selection by Cofea and the variant of removing PCA transformation. D, Memory use and running time for feature selection by Cofea and variant B of the parallel computing strategy. The x-coordinate is running time, and the y-coordinate is memory use. E, Cell clustering performance evaluated by NMI scores by noised-set features from with dropout rates selected by Cofea, HDA, epiScanpy and Signac. For each feature selection method, we set the dropout rate to be 0.05, 0.10, 0.15, 0.20, 0.25 and 0.30, and replaced random seeds 5 times for testing. The measure of center for the error bars and the dot on the line plot denote the mean value of NMI scores or overlapped proportion, and the error bar denotes the 95% confidence interval. F, Clustering results evaluated by NMI scores with different number of selected features from 3 TF-IDF transformations. G, Cell clustering performance evaluated by all six metrics (NMI, AMI, ARI, Homo, ASW and cLISI) with 20000 features selected from the PCA-transformed matrix. We set 10, 20, 50, 100, 150 and 200 as the number of PCs and tested 5 different random seeds. The dot on the line plot denotes the mean value of NMI scores, and the error bar denotes the 95% confidence interval. H, Cell clustering performance evaluated by all six metrics (NMI, AMI, ARI, Homo, ASW and cLISI) with 20000 features selected from different correlation coefficients processing.

#### Robustness to dropout events and hyperparameters

We further tested the robustness of Cofea when the two types of factors varied: (i) extrinsic factors, that is technical noise in the sequencing process; (ii) intrinsic factors, that is built-in hyperparameters in the framework of Cofea. We continued to use cell clustering on the Buenrostro2018 dataset to detect the impact of these two factors.

To mimic technical noise generated by sequencing platforms, we downsampled the reads in scCAS data with various dropout rates. More specifically, given a dropout rate, we randomly set the nonzero elements in the peak-by-cell count matrix to zero. We then compared the performance of Cofea with baseline methods using cell clustering. The results showed that Cofea consistently outperforms the baseline methods across the set of dropout rates from 5% to 30%. These findings indicate that Cofea is robust and can effectively handle data variations even in the presence of technical noise generated by the sequencing platform, suggesting that Cofea has great potential as a powerful tool for feature selection of scCAS data in a robust and reliable manner (Figure 5E). We also provided another perspective to test the performance of Cofea with technical noises (Supplementary Note S8 and Supplementary Figure S7).

To investigate the impact of hyperparameters, we varied each hyperparameter in Cofea in turn and fixed the other hyperparameters to the default setting. First, we noticed that although the TF-IDF transformation plays an important role in scCAS data analysis, the details of implementation vary among different toolkits. There are three most common implementations: (i) the original version of the TF-IDF transformation, widely used for scCAS data analysis (2,11,17,20,21,53-55), denoted as TF-IDF_original; (ii) improved TF-IDF modified by Signac(21), also as the default in Cofea, denoted as TF-IDF; (iii) another version proposed by scOpen (27), denoted as TF-IDF_scOpen (MATERIALS AND METHODS, and Supplementary Note S1). As shown in Figure 5F, Cofea with TF-IDF (the default setting in Cofea) gets better performance than the variants with other implementations of the TF-IDF transformation no matter how many features were required, which confirms the superiority of the TF-IDF in our method. Second, we varied the number of PCs. As shown in Figure 5G, with 10 to 200 PCs, Cofea achieves almost consistent performance, which demonstrates the robustness of Cofea to the number of PCs. Note that each PC may be potentially corresponding to a specific cell subpopulation, we thus set 100 as the default number of PCs to accommodate the analysis of large-scale datasets. Finally, we validated the contribution of PCC in comparison to alternative correlation coefficients, including SPCC and CSC (MATERIALS AND METHODS). As shown in Figure 5H, Cofea with PCC and CSC has almost consistent performance, slightly better than SPCC, indicating that Cofea is robust to the correlation coefficients. Different from PCC and CSC, SPCC only takes into account the rank of the elements rather than the values, leading to information loss, which may reason why SPCC got worse performance than the other two correlation coefficients. We also analysed the computational efficiency of Cofea (details in Supplementary Note S9 and Supplementary Figure S8). To summarize, Cofea achieves handle technical noise, adaptively selecting features on noised data, and has considerable robustness to the intrinsic factors, with stable performance on different built-in hyperparameters.

## Discussion

The high-noise and high-dimensional characteristics of scCAS data have increased the demand for a feature selection method, to accurately extract and preserve cellular heterogeneity when preprocessing scCAS data. To fill this gap, we developed a correlation-based feature selection method Cofea, for identifying biologically informative features in scCAS data. In contrast to the previous studies that focus on simplistic statistics of individual peaks, Cofea capitalizes on the intrinsical information from the peak-peak relationship. With a comprehensive evaluation on 54 published datasets with different sizes and qualities, we have demonstrated the advantages of Cofea in facilitating downstream analysis, particularly in dimensionality reduction and cell clustering. From the perspective of revealing biological signals, we illustrated that features selected by Cofea exhibit not only a significant overlap with cell type-specific accessible regions and candidate enhancers, but also encompass abundant chromatin regions related to biological functions and phenotypes. Through the quantitative assessment on simulated datasets, we showed the reasonable interpretation of Cofea’s superior performance. Furthermore, our results from ablation and dropout experiments also corroborated the elaborated design and high robustness of Cofea’s framework. Our experimental pipeline spans theoretical analysis, computational validation, performance evaluation, and biological interpretation, providing a novel research paradigm for the development of feature selection methods. Overall, Cofea offers an effective and valuable tool for scCAS data analysis, contributing to understanding cellular heterogeneity and detecting regulatory mechanisms.

Despite the progress achieved so far, we also have several potential directions for improving Cofea. First, considering that a substantial portion of scCAS data analytic tools are implemented in the R, such as Signac (21) and ArchR (56), we will develop an R version of Cofea, to offer researchers a richer pool of alternatives. Second, such a correlation-based framework can also be extended to scRNA-seq data or single-cell multi-omics data. It is interesting to consider how to incorporate the gene-gene or gene-peak intrinsic relationship with the feature selection step. Finally, we can leverage the intermediate outputs of Cofea to assess the purity of a cell population, as suggested by the recent study (57). In scRNA-seq data analysis, researchers commonly rely on marker genes to annotate cell types, while the limited availability of biomarkers in scCAS data poses challenges when interpreting results following cell clustering. Consequently, the assessment of whether cells from a given population have identical functions and state remains an urgent and formidable undertaking.

## DATA AVAILABILITY

The Buenrostro2018 dataset can be accessed from GEO under accession number GSE96772. The Joshua2021 dataset was collected from GEO with accession no. GSE166547. The Neurips2021 dataset can be accessed from GEO under accession number GSE194122. The Sai2020 dataset can be accessed from GEO with accession no. GSE140203. The MCA atlas is available under GEO with accession no. GSE111586. The HGCA atlas was collected from GEO with accession no. GSE184462. The FCA atlas is available at https://descartes.brotmanbaty.org/.

For CODE AVAILABILITY, The Cofea software, including detailed documents and tutorial, is freely available on GitHub (https://github.com/likeyi19/Cofea).

## SUPPLEMENTARY DATA

Supplementary Data are available at NAR online.

## AUTHOR CONTRIBUTIONS

R.J. and S.C. conceived the study and supervised the project. K.L. and X.C. designed, implemented and validated Cofea. S.S. and L.H. helped with analysing the results. X.C., K.L., R.J. and S.C. wrote the manuscript, with input from all the authors.

## ACKNOWLEDGEMENTS

We thank Yanhong Wu and Shirui Li for their helpful discussion and contributions to this research.

