## Supplementary Notes, Figures and Tables for "Cofea: correlation-based feature selection for single-cell chromatin accessibility data"

[1]Ministry of Education Key Laboratory of Bioinformatics, Bioinformatics Division at the Beijing National Research Center for Information Science and Technology, Center for Synthetic and Systems Biology, Department of Automation, Tsinghua University, Beijing 100084, China, [2]Center for Statistical Science, Department of Industrial Engineering, Tsinghua University, Beijing 100084, China, [3]Center for Statistical Science, Department of Industrial Engineering, Tsinghua University, Beijing, China, [4]School of Mathematical Sciences and LPMC, Nankai University, Tianjin 300071, China.

#These authors are equal contributors.

*To whom correspondence should be addressed.

Contact:

Contents

**Supplementary Notes** 3

**Supplementary Note S1: Alternative implementations of TF-IDF transformation** 3

**Supplementary Note S2: Tailored parallel arithmetic strategy** 4

**Supplementary Note S3: Alternative options for peak-peak correlation** 5

**Supplementary Note S4: Generation for simulated datasets** 6

**Supplementary Note S5: Metrics for assessment of dimensionality reduction and cell clustering** 8

**Supplementary Note S6: Detailed descriptions of the three baseline methods** 10

**Supplementary Note S7: Feature selection methods for scRNA-seq and their performance on scCAS data** 11

**Supplementary Note S8: Extended dropout experiments** 13

**Supplementary Note S9: Computational efficiency experiments** 14

**Supplementary Figures** 15

**Supplementary Figure S1** 15

**Supplementary Figure S2** 17

**Supplementary Figure S3** 19

**Supplementary Figure S4** 21

**Supplementary Figure S5** 22

**Supplementary Figure S6** 23

**Supplementary Figure S7** 24

**Supplementary Figure S8** 25

**Supplementary Tables** 26

**Supplementary Table S1 Real datasets information summary** 26

**References** 29

### Supplementary Notes

#### Supplementary Note S1: Alternative implementations of TF-IDF transformation

We adopt two other implementations of TF-IDF transformation from its original version (referred to as TF-IDF_original transformation) and scOpen[1] improved version (referred to as TF-IDF_scOpen transformation). The three implementations of TF-IDF transformation serves as a hyperparameter for Cofea, which is presented as an optional choice within the model. The TF-IDF_original transformation is defined as:

$$x''_{ij} = \frac{x_{ij}}{\sum_i x_{ij}} \cdot \log\left(1 + \frac{n}{\sum_j x_{ij}}\right)$$

where $x_{ij}$ denotes the element in the peak-by-cell matrix $\boldsymbol{X} \in \mathbb{R}^{p \times n}$, and $x''_{ij}$ is the corresponding element in the newly formed matrix $\boldsymbol{X}'' \in \mathbb{R}^{p \times n}$ after TF-IDF_original transformation.

The TF-IDF_scOpen transformation uses a characteristic function $\mathrm{I}(x_{ij} > 0)$ to represent whether peak $i$ is open or closed in cell $j$:

$$\mathrm{I}(x_{ij} > 0) = \begin{cases} 1, & x_{ij} > 0 \\ 0, & x_{ij} = 0 \end{cases}$$

The TF-IDF_scOpen transformation can be calculated based on this representation:

$$x'''_{ij} = \frac{\mathrm{I}(x_{ij} > 0)}{\sum_i \mathrm{I}(x_{ij} > 0)} \cdot \log\left(\frac{n}{\sum_j \mathrm{I}(x_{ij} > 0)}\right)$$

where $x'''_{ij}$ is the corresponding element in the newly formed matrix $\boldsymbol{X}''' \in \mathbb{R}^{p \times n}$ after TF-IDF_scOpen transformation. We utilized the implementation presented in the scOpen source code, specifically the TfidfTransformer model from scikit-learn package with the same parameter settings as in scOpen.

**Supplementary Note S2: Tailored parallel arithmetic strategy**

As Cofea is a correlation-based feature selection method, obtaining correlation coefficients between peaks is an essential step in its procedure. However, as the number of peaks typically reaches 100,000, the peak-by-peak correlation matrix is exceedingly high dimensional, and the elements within it are continuous and dense. This will raise issues such as memory crush or intolerable running time. Moreover, as mentioned in the framework of Cofea, only the mean value and mean square value of inter-peak correlation coefficients are required, rather than the whole peak-by-peak correlation matrix. If the element $c_{ij}$ in $\boldsymbol{C}$ is computed iteratively, that is, for each peak, an inner loop is executed to obtain Pearson correlation coefficients (PCC) between that peak and the other peaks, and then store the mean value and mean square value. This process is repeated $p$ times, using an outer loop to iterate over each peak, which will take up plenty of time, causing low computing efficiency. Therefore, we tailored a parallel arithmetic strategy to obtain the correlation between peaks, computing PCC between $k$ (specified by the user) peaks and all peaks at one time. More specifically, we used matrix operations to accelerate the computation. We took out a submatrix from the PCA-transformed matrix $\boldsymbol{P}$, referred to as $\boldsymbol{P}^{(i)} \in \mathbb{R}^{k\times q}$, containing the $i$th to $(i+k)$th peaks and all cells. Then we extended the formula of PCC into matrix operation and calculated the peak-peak correlation matrix $\boldsymbol{C}_i \in \mathbb{R}^{k\times p}$, which contains elements from $c_{i1}$ to $c_{(i+k)p}$ in $\boldsymbol{C}$.

We first calculated a vector $\boldsymbol{v} \in \mathbb{R}^p$ using the formula:

$$v_m = \frac{1}{n}\sum_{j}^{n}\left(p_{mj} - \frac{1}{n}\sum_{i}^{n} p_{mi}\right)^2$$

where $v_m$ is an element of $\boldsymbol{v}$, representing the variance of the accessibility of peak $m$ across all cells. A matrix $\boldsymbol{V} \in \mathbb{R}^{k\times p}$ is generated from $\boldsymbol{v}$ by vertically replicating $k$ times. Next, we calculate the correlation matrix as:

$$\boldsymbol{C}_i = \frac{\left(\boldsymbol{P}^{(i)} - \overline{\boldsymbol{P}}^{(i)}\right) \times (\boldsymbol{P} - \overline{\boldsymbol{P}}).\mathrm{T}}{\sqrt{\mathrm{Diag}\left((\boldsymbol{P}^{(i)} - \overline{\boldsymbol{P}}^{(i)}) \times (\boldsymbol{P}^{(i)} - \overline{\boldsymbol{P}}^{(i)}).\mathrm{T}\right)} \times \sqrt{n\boldsymbol{V}}}$$

where $\overline{\boldsymbol{P}} \in \mathbb{R}^{p\times q}$ and $\overline{\boldsymbol{P}}^{(i)} \in \mathbb{R}^{k\times q}$ are matrices that element denotes the mean value of the corresponding row in $\boldsymbol{P}$ and $\boldsymbol{P}^{(i)}$, therefore the elements in the same row share a consistent value. $\mathrm{Diag}(\boldsymbol{A})$ denotes a function that keeps only the diagonal elements of $\boldsymbol{A}$ and sets other elements to 0.

**Supplementary Note S3: Alternative options for peak-peak correlation**

Besides Pearson correlation coefficients (PCC), we also provided Spearman correlation coefficients (SPCC) and Cosine similarity coefficients (CSC) as alternatives. The method to obtain the inter-peak correlation cofficients also serves as an optional hyperparameter for Cofea. SPCC can be calculated as the formula:

$$SPCC(\boldsymbol{p}_{k\cdot},\boldsymbol{p}_{l\cdot}) = 1 - \frac{6\sum_j^n \left(\mathrm{R}(p_{kj}) - \mathrm{R}(p_{lj})\right)^2}{n(n^2-1)}$$

where $\boldsymbol{p}_{k\cdot}$ represents the $k$th peak, while $p_{kj}$ represents the element of PCA-transformed matrix $\boldsymbol{P}$, and $n$ is the number of cells. The function $\mathrm{R}(p_{kj})$ denotes the rank of the $j$th element in vector $\boldsymbol{p}_{k\cdot}$.

CSC uses the cosine of the angle between two vectors to measure the similarity, and can be formulated as:

$$CSC(\boldsymbol{p}_{k\cdot},\boldsymbol{p}_{l\cdot}) = \frac{\sum_j^n p_{kj}\cdot p_{lj}}{\sqrt{\sum_j^n p_{kj}^2}\sqrt{\sum_j^n p_{lj}^2}}$$

**Supplementary Note S4: Generation for simulated datasets**

For dataset S1, we generated a peak-by-cell matrix with 100,000 peaks and 2,000 cells. S1 contains two cell types: cell type A (1000 cells) and cell type B (1000 cells), and for each cell type, cell type-specific peaks account for 5% of all peaks. For background peaks, we set the degree of accessibility to 5%, that is, for each corresponding row vector, we randomly the value of 5% elements to 1 and keep remaining elements to 0. To simulate cell type-specific peaks, we set the degree of accessibility from 5% to 10% with a uniform distribution in one cell type, and simultaneously set the degree of accessibility from 5% to 0% in another cell type. By performing feature selection on dataset S1, we verify whether Cofea is capable of identifying cell type-specific peaks that have consistent degree of accessibility consistent with that of background peaks. To further test performance when the accessibility of peaks varies, we changed the degree of accessibility from a fixed value to a dynamic range to simulate dataset S2. In dataset S2, the degree of background peak accessibility ranges from 0% to 5%, while for cell type-specific peaks, we set the degree of accessibility ranging from 0% to 7.5% in one cell type, and simultaneously set the degree in the other cell type from 0% to 2.5%. Other settings in the generation procedure of dataset S2 are consistent with those in S1.

To generate the real peak-by-cell matrix, peak calling is a fundamental step for captures the highest accessibility signals from genomic regions (peaks). However, aggregating accessibility signals from abundant single cells is required to identify such peaks, and inevitably, detecting the biological signals from the rare cell type is challenging because of its small proportion. We hereby generated dataset S3 and S4, for evaluating performance of Cofea when applied to imbalanced cell populations. S3 contains 100,000 peaks and 3000 cells in three cell types: cell type A (1500 cells), cell type B (750 cells) and cell type C (750 cells). For each cell type, cell type-specific peaks account for 5% of all peaks. For background peaks, we set the degree of accessibility ranging from 0% to 5% with a uniform distribution across all cells. For cell type-specific peaks, we set the degree of accessibility ranging from 5% to 10% in one cell type and from 1% to 3% in the other cell types. S4 contains 100,000 peaks and 4000 cells, including three common cell types (cell type A, B and C) with 1,200 cells within each cell type, and a rare cell type D with 400 cells. For each cell type, cell type-specific peaks account for 5% of all peaks. For background peaks, we set the degree of accessibility ranging from 0% to 5% with a uniformly distribution across all cells. The degree of accessibility for cell type-specific peaks and background peaks were set the same as S3.

In comparison to dataset S1 to S4 with discrete cell populations, dataset S5 is constituted by 750 cells with a continuous differentiation trajectory, that is, 250 cells of cell type A differentiating into 250 cells of cell type B and 250 cells of cell type C. Using S5 for illustration, we tested whether features identified by Cofea could be effectively implemented in trajectory inference. As a side note, here the cell type-specific peaks are no longer more accessible in a tailored cell type, but relevant to cell differentiation. We set the number of peaks to 100,000 and treated 10% of all peaks as cell type-specific peaks. To reflect cell differentiation from the accessibility of peaks, we generate the value of $i$th peak and $j$th cell from a Bernoulli distribution, of which the probability parameter $\theta_{ij}$ follows a Brownian motion. More specifically, for each peak, the increment $\varepsilon$ between $j$th cell and $j+1$th cell is sampled from a Gaussian distribution $\mathcal{N}(0,0.0001)$, that is $\theta_{i(j+1)} = \theta_{ij} + \varepsilon$, and for the root cell, $\theta_{i1}$ are fixed to 0.05. Note that to generate high-fidelity data, the parameter $\theta_{ij}$ is restricted to a range of $[0,0.2]$. The accessibility of background peaks is expected to be consistent across different cell types. To differentiate between cell type-specific peaks and background peaks, we randomly shuffled the accessibility of each background peak in cells while maintaining the range of 0 and 0.2.

**Supplementary Note S5: Metrics for assessment of dimensionality reduction and cell clustering**

We denote $\boldsymbol{U}$ as the ground-truth cell labels, $\boldsymbol{V}$ as the predicted cluster labels, and the NMI score can be calculated by the following formula:

$$NMI = \frac{MI(\boldsymbol{U},\boldsymbol{V})}{\sqrt{H(\boldsymbol{U}) \cdot H(\boldsymbol{V})}}$$

where $\mathrm{MI}(\cdot,\cdot)$ is used to compute the mutual entropy, and $\mathrm{H}(\cdot)$ is used to compute the entropy.

The AMI score can be calculated as follows:

$$AMI = \frac{MI(\boldsymbol{U},\boldsymbol{V}) - E\big(MI(\boldsymbol{U},\boldsymbol{V})\big)}{avg\big(H(\boldsymbol{U}),H(\boldsymbol{V})\big) - E\big(MI(\boldsymbol{U},\boldsymbol{V})\big)}$$

where $\mathrm{E}(\cdot)$ is the expectation function. We then suppose that $u_i$ is the $i$th cell label, $v_j$ is the $j$th cluster label, $n_{ij}$ is the number of cells simultaneously belonging to $u_i$ and $v_j$, $n_{i\cdot}$ is the number of cells that belong to $u_i$, $n_{\cdot j}$ is the number of cells that belong to $v_j$, and $n$ is the total number of cells. The ARI score is calculated as follows:

$$ARI = \frac{\sum_{i,j}\binom{n_{ij}}{2} - \left[\sum_i \binom{n_{i\cdot}}{2}\sum_j \binom{n_{\cdot j}}{2}\right]/\binom{n}{2}}{\frac{1}{2}\left[\sum_i \binom{n_{i\cdot}}{2} + \sum_j \binom{n_{\cdot j}}{2}\right] - \left[\sum_i \binom{n_{i\cdot}}{2}\sum_j \binom{n_{\cdot j}}{2}\right]/\binom{n}{2}}$$

The Homo score can be calculated by the following formula:

$$Homo = 1 - \frac{H(\boldsymbol{U}|\boldsymbol{V})}{H(\boldsymbol{U})}$$

where $\mathrm{H}(\boldsymbol{U}|\boldsymbol{V})$ is the entropy of the ground-truth cell labels based on the predicted cluster labels. For all the four metrics, a higher score closed to 1 indicates that the clustering results are more consistent with the ground-truth labels.

The silhouette width (SW) measures the relationship between the distances of a cell from other cells in the same cluster (referred to as inner-cluster distances) and the distances of the cell from cells in the closest cluster (referred to as inter-cluster distances). The SW score for a single cell is computed as follows:

$$SW = \frac{b - a}{max(a,b)}$$

where $a$ is the average of inner-cluster distances and $b$ is the average of inter-cluster distances. ASW is the average of all the silhouette widths of a set of cells, assessing whether the clusters are dense and well-separated, ranging between -1 and 1. cLISI is used to measure the accuracy of an embedding with cell-type prediction. Specifically, after mixing the cells, cLISI predict type for each cell based on its 30 nearest neighbors in low-dimensional space, with the weighted sum of their cell types to find out the most likely prediction. A cLISI of 1 reflects successful separation of cell types, that is, different cell types group separately instead of together.

**Supplementary Note S6: Detailed descriptions of the three baseline methods**

Selecting peaks with highest degree of accessibility (HDA) is the simplest and most widely used feature selection method in scCAS data analysis[2, 3]. The HDA algorithm aggregates the read counts of each feature across all single-cells, and ranks the features based on this cumulative count. The features with the highest degree of accessibility are selected as the "informative features".

epiScanpy[4] is one of the most commonly used toolkits for scCAS data analysis in Python language, and feature selection is a step in its preprocessing pipeline. epiScanpy considers the most variable features to open only in half of the cells, and the least variable features to open in none or all of the cells. To this end, epiScanpy defines the variability score (VS) of a feature as the following formula:

$$VS_i = 1 - \left| \sum_{j}^{n} x_{ij} - 0.5 \right|$$

where $x_{ij}$ denotes the element in the raw peak-by-cell count matrix $\boldsymbol{X} \in \mathbb{R}^{p \times n}$. epiScanpy selects a user-defined number of features with the highest variability score as the "informative features".

Signac[5] is one of the most widely used toolkit for scCAS data analysis in R language. In its pipeline, feature selection is performed after peak calling and cell filtering. Signac performs TF-IDF transformation on the peak-by-cell matrix, followed by computing the quantile of the sum of each row in the transformed matrix. The quantile indicates the importance score of each feature, and features with higher quantiles are considered as the "informative features".

**Supplementary Note S7: Feature selection methods for scRNA-seq and their performance on scCAS data**

Many computational methods have been proposed for feature selection on scRNA-seq data, namely HVG, M3drop, NBdrop, and GiniClust. We compared these four methods and tested its performance on scCAS data. HVG suggests that genes with coefficients of variation (CV) across cells are more informative. The CV of each gene is defined as:

$$CV_i^2 = \frac{\sigma_i^2}{\mu_i^2}$$

where $\mu_i$ and $\sigma_i^2$ denote the mean and variance value of the expression of gene $i$ across all cells. HVG fits the relationship between $CV$ and $\mu$:

$$CV^2 = \alpha_0 + \frac{a_1}{\mu}$$

where $\alpha_0$ and $a_1$ are the two parameters obtained by model fitting. Statistical test is used to test the variability of each gene, and features with higher residual are selected.

M3drop uses the Michaelis-Menten function to model the relationship between mean expression and dropout rate value of gene expression. To assess the significance of genes, a t-test is conducted to determine if the gene-specific parameter $K_i$ is equal to the common parameter $K_M$ in the Michaelis-Menten function. NBdrop treats the gene expression matrix as a negative binomial distribution, and dropout rate of each gene is estimated by the parameters of this distribution model. NBdrop selects genes based on the estimated dropout rates. GiniClust fits the relationship between the gini coefficient and the maximum gene expression to select genes. We tested these four methods on a scRNA-seq dataset of Peripheral blood monoculear cells (PBMC), and the results are shown in the left part of Supplementary Figure S1.

The above methods share a common characteristic in that they all fit a model to represent the relationship between two statistics of gene expression, and identify genes based on the residual of the fitted model. However, as shown in Supplementary Figure S1, upon applying these approaches to a scCAS dataset, namely the MCA kidney dataset, we have observed non-conformance with their intended design. Specifically, the primary objective of these methods is to identify significant features by selecting outliers of the fitted curve, but within each method, there is a lack of outliers and instead conformed to a smooth curve, indicating the invalid assumption on scCAS data. An error was encountered while calculating the Gini coefficient due to the binary characteristic of the data. In summary, feature selection methods specifically designed for scRNA-seq data have evident errors and limitations when applied to scCAS data, and consequently, we did not continue to serve these methods as baseline method for comprehensive benchmarking.

**Supplementary Note S8: Extended dropout experiments**

To verify whether Cofea is able to adapt to the technical noise, we downsampled the reads on the Buenrostro2018 dataset with various dropout rates. More specifically, given a dropout rate, we randomly set the nonzero elements in the peak-by-cell count matrix to zero. We compared the overlapped proportion of selected features on raw data (referred as to raw-set features) with features selected on noised data (referred as to noised-set features). Interestingly, we observed a trend when incorporating more noise with count matrix, features selected by Cofea have a smaller overlap with raw-set features, while baseline methods basically remain the original selection (Supplementary Figure S7A). To validate if such the change of Cofea is beneficial for downstream analysis, we performed dimensionality reduction and cell clustering on the noised data with raw-set features and noised-set features, respectively. As shown in Supplementary S7B, Cofea with noised-set features achieved a relatively better performance, suggesting that the feature selected by Cofea are adaptable to technical noise.

**Supplementary Note S9: Computational efficiency experiments**

We assessed the computational efficiency and scalability of Cofea using the HGCA esophagus dataset, which contains 82469 cells and 101251 peaks after preprocessing. We randomly downsampled cells or peaks to obtain a variety of newly-formed datasets with different sizes, and then benchmarked the running time and peak memory usage of different feature selection methods. Note that since Cofea and baseline methods evaluates and ranks all the peaks regardless of how many features users require, the number of selected features has no effect on the evaluation. As shown in Supplementary Figure S8A, the running time and memory use of Cofea increase as the number of cells or peaks increases. More specifically, when the dataset contains more than 50,000 cells, running time required for the stepwise preprocessing takes up the majority of the total running time and was proportional to the size of the dataset. The running time taken for correlation calculation and fitting steps, on the other hand, is hardly affected by the number of cells due to the implementation of cell-wise PCA steps (Supplementary Figure S8B). We also compared Cofea with other baseline methods on the complete HGCA heart dataset, and it is intuitive that, because of the need to obtaining inter-peak correlation in Cofea, the computational expense is larger than methods based on simplistic statistics of individual peaks (Supplementary Figure S8C).

### Supplementary Figures

#### Supplementary Figure S1

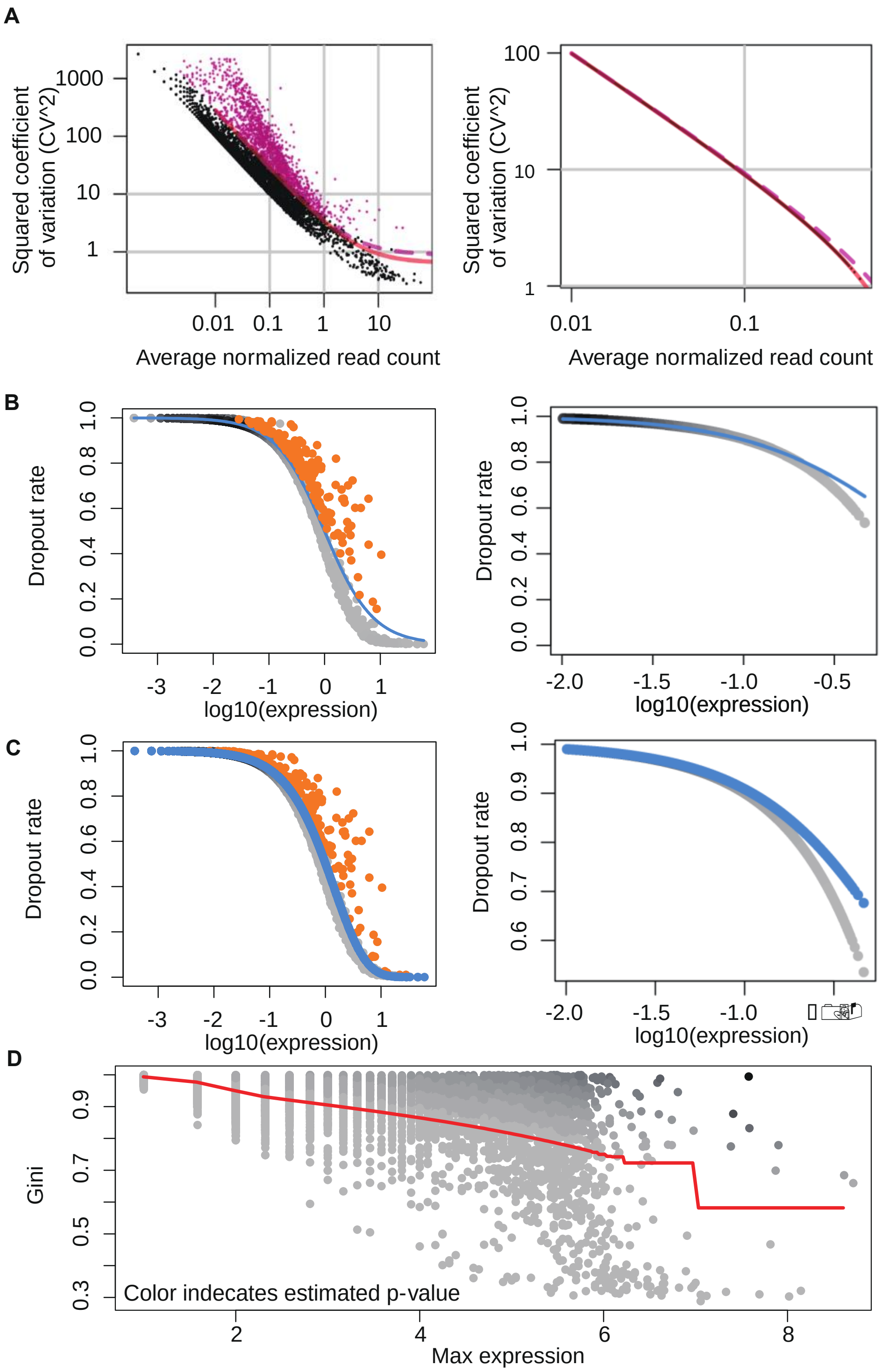


**Supplementary Figure S1.** The application of feature selection method for scRNA-seq data to scRNA-seq data and scCAS data. Perform feature selection using (A) HVG, (B) M3Drop, (C) NBDrop on the Peripheral blood monoculear cell (PBMC) and MCA Kidney datasets. PBMC dataset is a scRNA-seq dataset and arrayed on the left, while MCA Kidney is a scCAS dataset and arrayed on the right. D, Feature selection on the PBMC dataset using GiniClust. Due to the binarized nature of the scCAS data, GiniClust throws an error during the computation process, so the corresponding results are not provided here. The points in the figure are representations of features under specific two-dimensional characteristics, and the lines are obtained according to the fit of all genes. The highlighted points (pink points in A and orange points in B and C, respectively) are the informative features that have been selected, while the black and grey points serve as non-informative features.

### Supplementary Figure S2

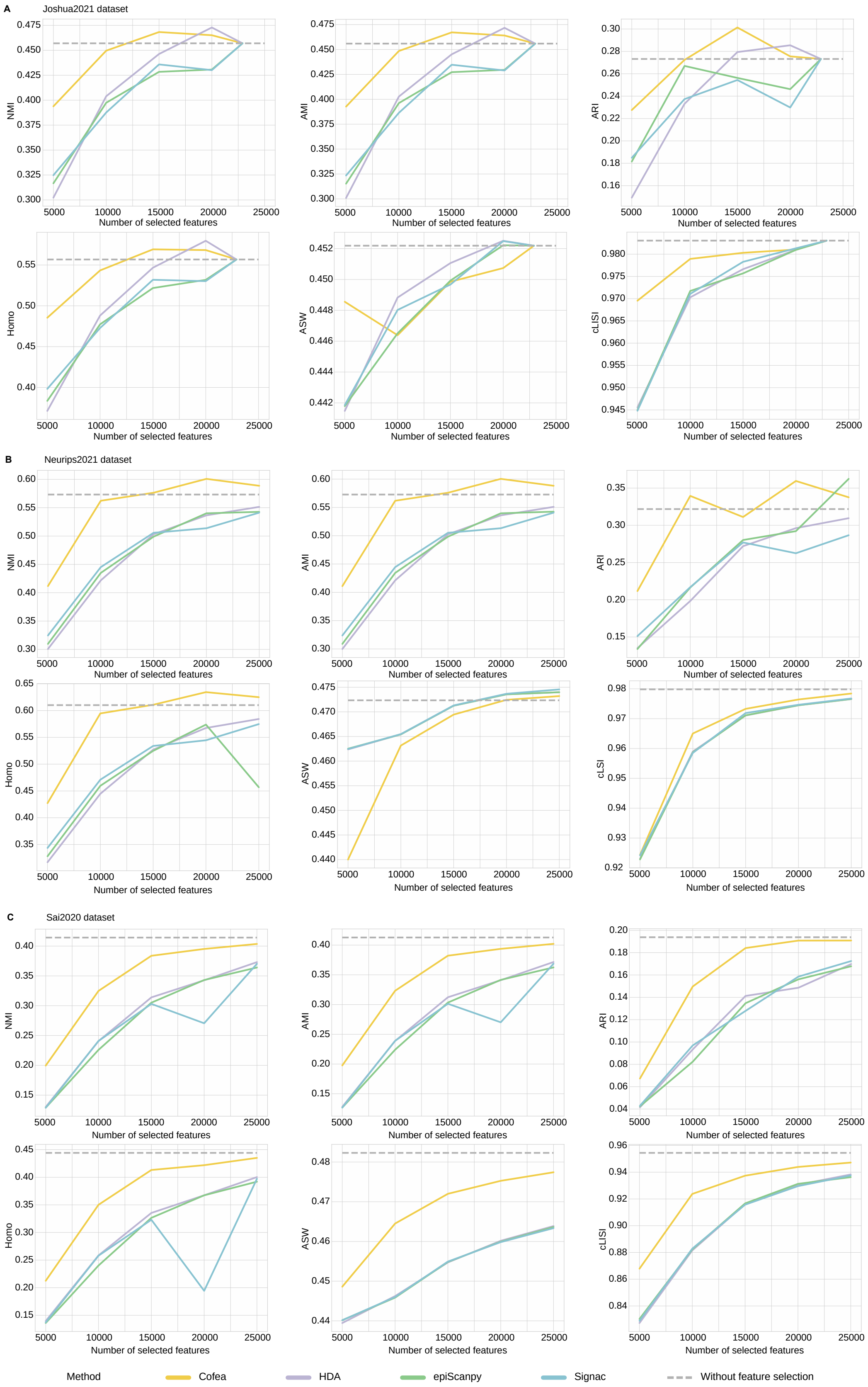


**Supplementary Figure S2.** Evaluation with various number of selected features on the Joshua2021,

Neurips2021 and Sai2020 datasets using NMI, ARI, AMI, Homo, ASW and cLISI scores. A, Dimensionality reduction and cell clustering performance on the Joshua2021 dataset, using features selected by Cofea, HDA, epiScanpy and Signac, respectively. For each feature selection method, we set 5,000, 10,000, 15,000, 20,000 and 25,000 as the number of selected features. B-D, Perform the same operations on the (B) Neurips2021 dataset, (C) Wang2022 dataset and (D) Sai2020 dataset, respectively.

**Supplementary Figure S3**

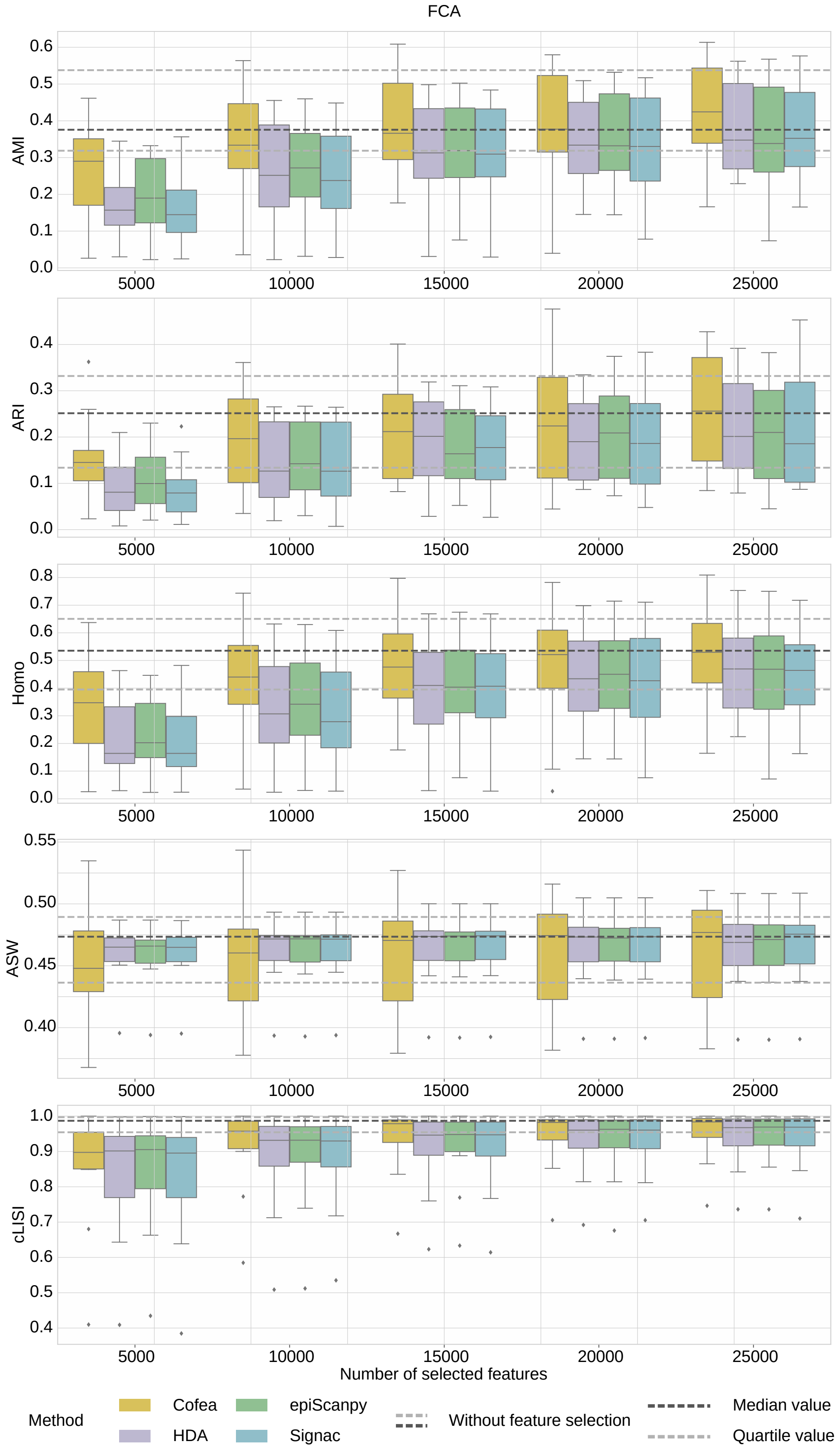


**Supplementary Figure S3.** Evaluation using the remaining five metrics with various numbers of selected features on datasets in FCA atlas. Cell clustering performance evaluated by AMI, ARI,

Homo, ASW and cLISI scores using different number of features selected by Cofea, HDA, epiScanpy and Signac, respectively, on datasets in FCA atlas. For each feature selection method, we set 5000, 10000, 15000, 20000 and 25000 as the number of selected features. The measure of center for the error bars denotes the median value of the metric on different datasets in an atlas, and the error bar denotes the maximum and minimum value after removing the outliers, which are defined as the values whose distance from the median is greater than 1.5 times the quartile distance. The black dotted line denotes the median score of cell clustering without feature selection, and the gray dotted lines denote the first and third quartiles.

### Supplementary Figure S4

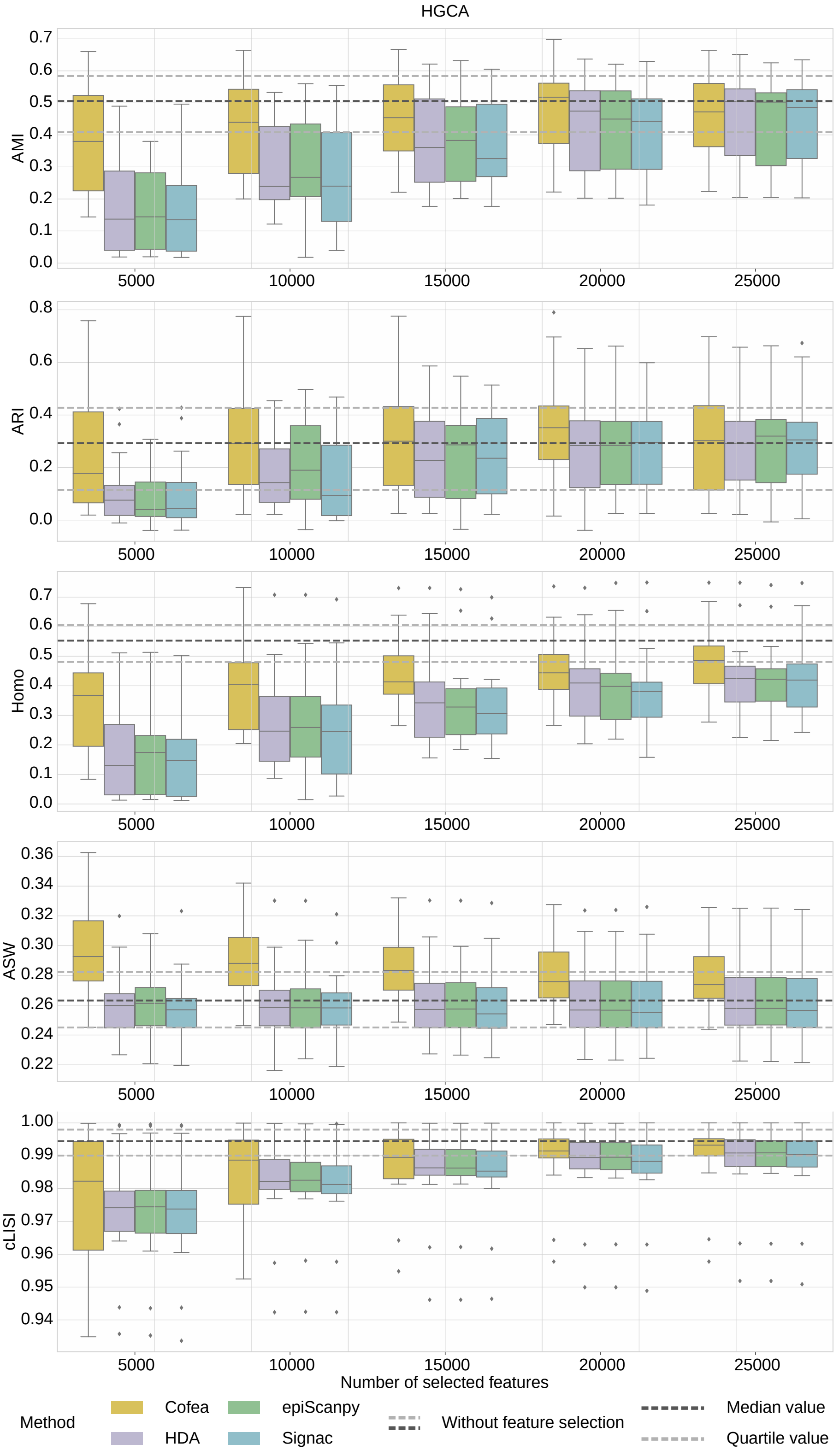


**Supplementary Figure S4.** Evaluation using the remaining five metrics with various number of selected features on datasets in HGCA atlas.

### Supplementary Figure S5

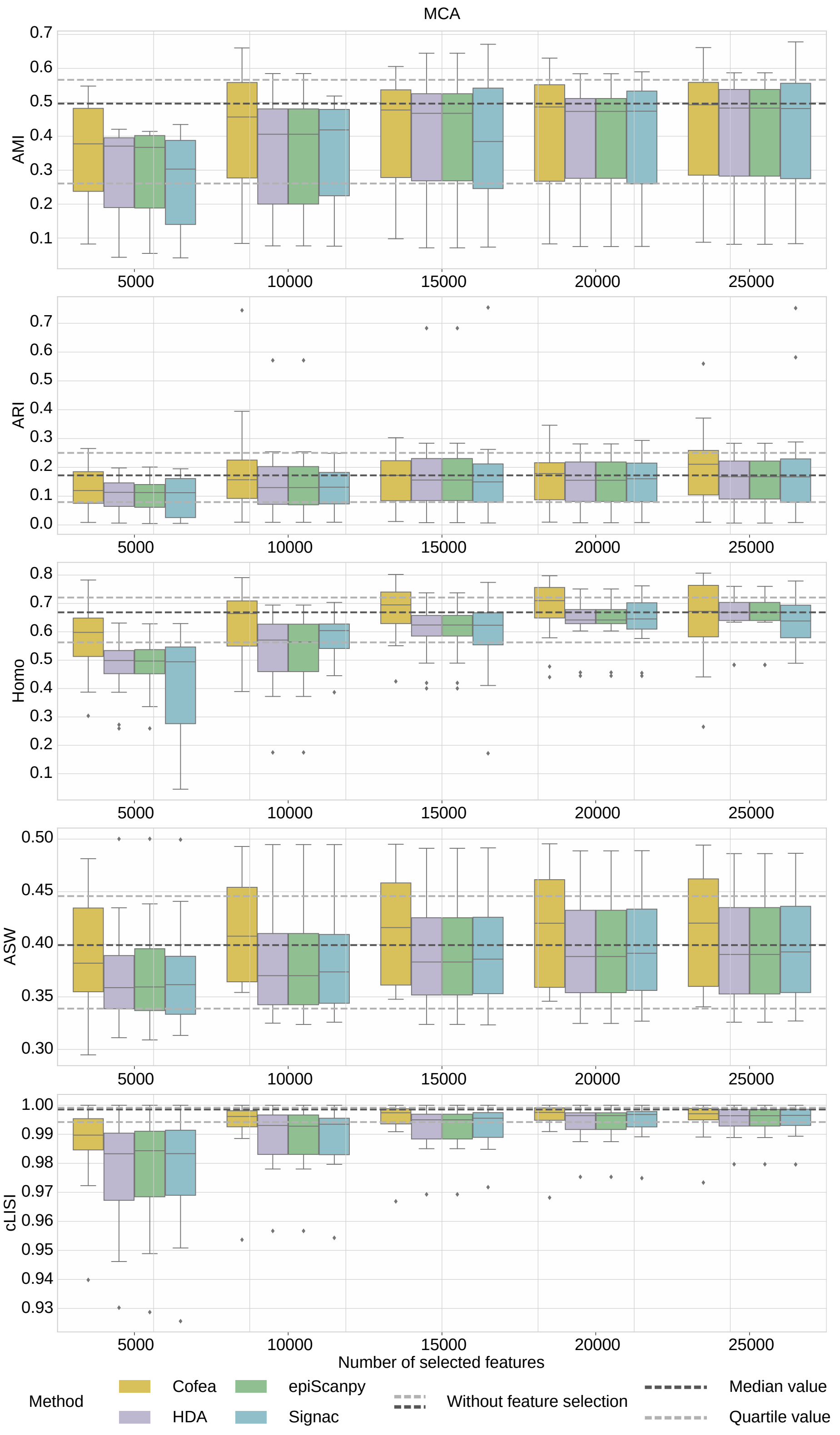


**Supplementary Figure S5.** Evaluation using the remaining five metrics with various number of selected features on datasets in MCA atlas.

### Supplementary Figure S6

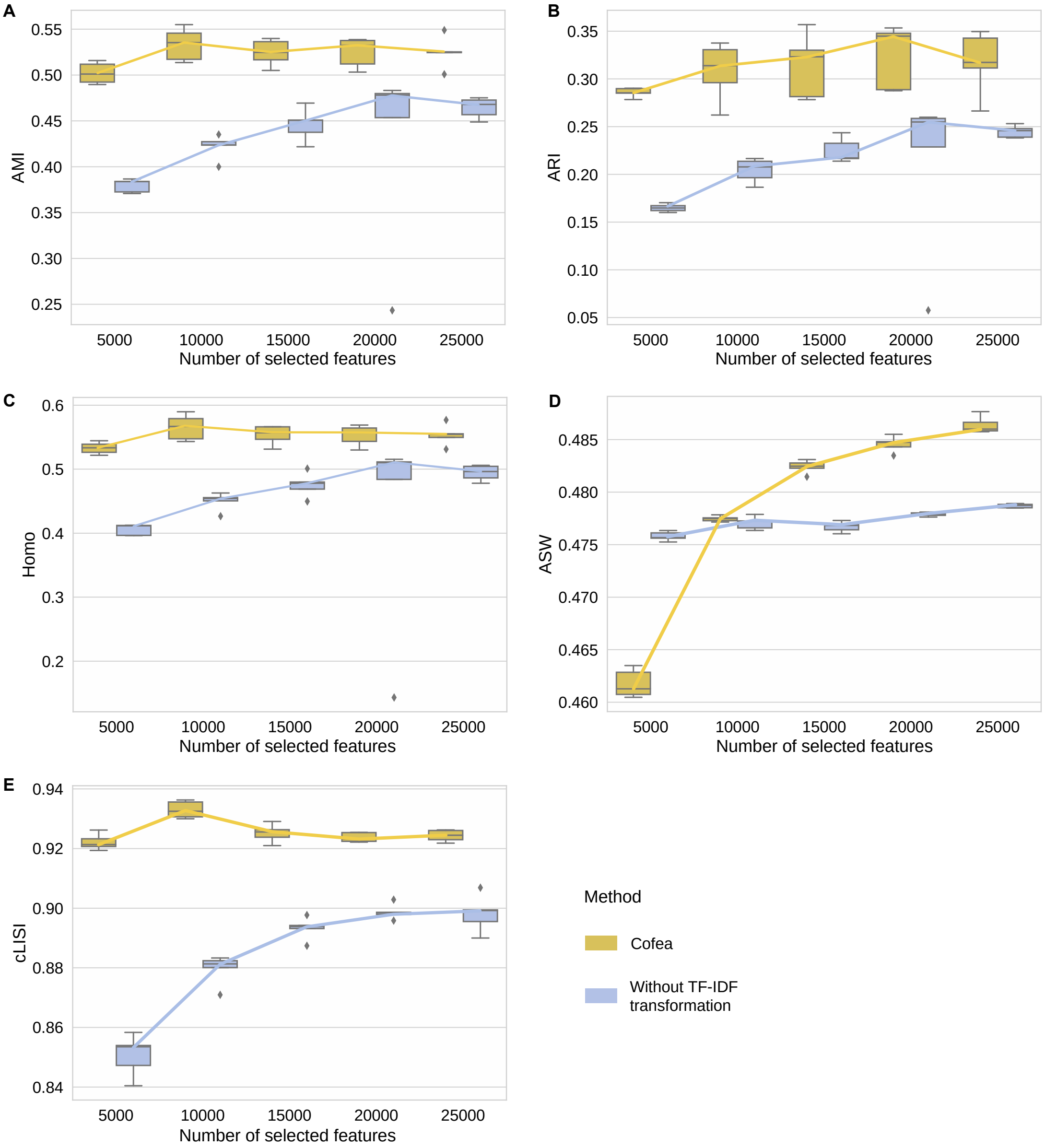


**Supplementary Figure S6.** Cell clustering performance evaluated by the remaining five metrics, that is, (A) AMI, (B) ARI, (C) Homo, (D) ASW, (E) cLISI, with different number of selected features from Cofea and the variant by removing TF-IDF transformation. The measure of center for the error bars denotes the median value, and the error bar denotes the maximum and minimum value after removing the outliers, which are defined as the values whose distance from the median is greater than 1.5 times the quartile distance.

### Supplementary Figure S7

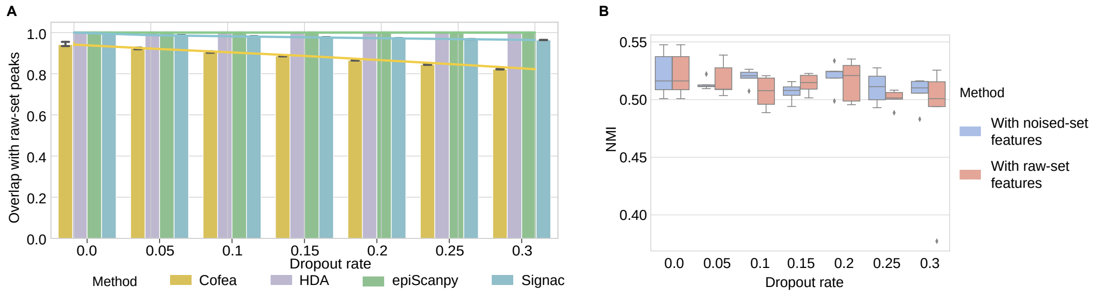


**Supplementary Figure S7.** Cofea can be adaptable to the technical noise. A, Overlapped proportion with raw-set peaks from dropout rates selected by Cofea, HDA, epiScanpy and Signac. For each feature selection method, we set the dropout rate to be 0.05, 0.10, 0.15, 0.20, 0.25 and 0.30, and replaced random seeds 5 times for testing. The measure of center for the error bars and the dot on the line plot denote the mean value of NMI scores or overlapped proportion, and the error bar denotes the 95% confidence interval. B, Clustering results evaluated by NMI scores with noised-set and raw-set features selected from datasets with different dropout rates. The measure of center for the error bars denotes the median value of NMI scores, and the error bar denotes the maximum and minimum value after removing the outliers, which are defined as the values whose distance from the median is greater than 1.5 times the quartile distance.

**Supplementary Figure S8**

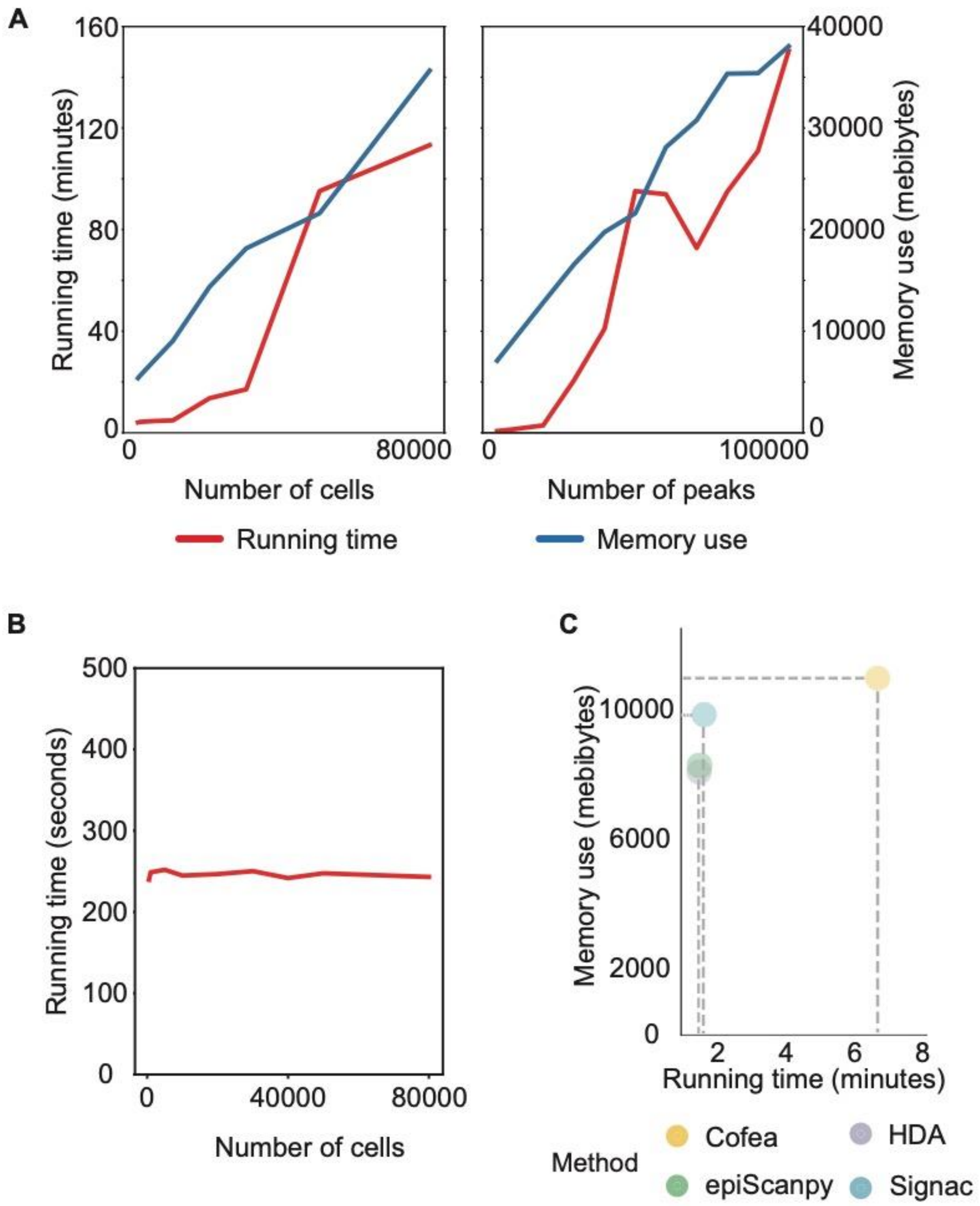


**Supplementary Figure S8.** Computational efficiency analysis using the HGCA esophagus dataset. A, Running time and memory usage of Cofea with number of cells and peaks. B, Running time of Cofea with different number of cells. C, Memory use and running time of Cofea, HDA, epiScanpy, and Signac, respectively.

### Supplementary Tables

#### Supplementary Table S1 Real datasets information summary

The cell number in the table is the number of sequenced cells contained in the dataset. Peak number (pre-filtration) refers to the number of accessible regions of chromatin detected, and peak number (after filtration) refers to the number of peak remaining after the selection to be accessible in more than 1% of cells. Binary is used to indicate whether a dataset is binary, and Sparsity refers to the proportion of 0 counts in the peak-cell matrix.

| Dataset | Cell number | Peak number (pre-filtration) | Peak number (after filtration) | Binary | Sparsity |
|---|---|---|---|---|---|
| Buenrostro2018 | 2,034 | 455,057 | 100,934 | False | 98.972% |
| Joshua2021 | 15,298 | 91,754 | 22,909 | False | 98.675% |
| Neurips2021 | 69,249 | 116,490 | 65,336 | True | 96.921% |
| Sai2020 | 34,774 | 344,592 | 97,669 | False | 98.834% |
| MCA bone marrow | 8,403 | 436,177 | 86,856 | True | 99.153% |
| MCA cerebellum | 2,278 | 436,177 | 73,012 | True | 99.178% |
| MCA heart | 7,650 | 436,177 | 95,491 | True | 98.927% |
| MCA intestine | 11,163 | 436,177 | 110,762 | True | 99.078% |
| MCA kidney | 6,431 | 436,177 | 110,928 | True | 98.688% |
| MCA large intestine | 7,086 | 436,177 | 114,751 | True | 99.018% |
| MCA liver | 6,167 | 436,177 | 100,499 | True | 98.612% |
| MCA lung | 9,996 | 436,177 | 72,598 | True | 99.139% |
| MCA prefrontal cortex | 5,959 | 436,177 | 190,299 | True | 98.010% |
| MCA small intestine | 4,077 | 436,177 | 102,660 | True | 99.181% |
| MCA spleen | 4,020 | 436,177 | 79,819 | True | 98.737% |
| MCA testis | 2,723 | 436,177 | 143,302 | True | 98.936% |

| | | | | | |
|---|---|---|---|---|---|
| MCA thymus | 7,617 | 436,177 | 86,568 | True | 98.649% |
| MCA whole brain | 8,766 | 436,177 | 160,516 | True | 98.458% |
| FCA adrenal | 63,958 | 1,050,819 | 70,522 | False | 99.601% |
| FCA cerebellum | 3,313 | 1,050,819 | 89,256 | False | 99.500% |
| FCA cerebrum | 85,261 | 1,050,819 | 23,954 | False | 99.828% |
| FCA eye | 9,712 | 1,050,819 | 126,193 | False | 99.296% |
| FCA heart | 79,248 | 1,050,819 | 99,607 | False | 99.447% |
| FCA intestine | 42,942 | 1,050,819 | 64,647 | False | 99.619% |
| FCA kidney | 75,490 | 1,050,819 | 126,193 | False | 99.464% |
| FCA liver | 183,175 | 1,050,819 | 22,092 | False | 99.823% |
| FCA lung | 72,662 | 1,050,819 | 58,521 | False | 99.658% |
| FCA muscle | 27,181 | 1,050,819 | 96,720 | False | 99.447% |
| FCA pancreas | 4,994 | 1,050,819 | 111,039 | False | 99.419% |
| FCA placenta | 45,181 | 1,050,819 | 85,540 | False | 99.505% |
| FCA spleen | 2,157 | 1,050,819 | 40,187 | False | 99.729% |
| FCA stomach | 3,840 | 1,050,819 | 126,706 | False | 99.348% |
| FCA thymus | 21,499 | 1,050,819 | 59,007 | False | 99.574% |
| HGCA adipose | 16,243 | 1,154,611 | 64,880 | False | 99.603% |
| HGCA artery | 46,283 | 1,154,611 | 158,344 | False | 99.295% |
| HGCA colon | 55,084 | 1,154,611 | 70,845 | False | 99.574% |
| HGCA heart | 111,773 | 1,154,611 | 87,544 | False | 99.516% |
| HGCA liver | 6,498 | 1,154,611 | 85,269 | False | 99.537% |
| HGCA mammary | 16,410 | 1,154,611 | 31,223 | False | 99.783% |
| HGCA nerve | 13,866 | 1,154,611 | 95,925 | False | 99.482% |
| HGCA pancreas | 33,221 | 1,154,611 | 56,427 | False | 99.611% |
| HGCA small intestine | 17,788 | 1,154,611 | 52,270 | False | 99.685% |
| HGCA thyroid | 28,299 | 1,154,611 | 40,066 | False | 99.710% |
| HGCA vagina | 4,824 | 1,154,611 | 94,173 | False | 99.534% |

| | | | | | |
|---|---|---|---|---|---|
| HGCA adrenal | 7,796 | 1,154,611 | 39,614 | False | 99.734% |
| HGCA brain | 14,906 | 1,154,611 | 54,794 | False | 99.672% |
| HGCA esophagus | 82,469 | 1,154,611 | 101,251 | False | 99.469% |
| HGCA islet | 14,298 | 1,154,611 | 126.021 | False | 99.408% |
| HGCA lung | 41,349 | 1,154,611 | 73,638 | False | 99.601% |
| HGCA muscle | 43,553 | 1,154,611 | 53,357 | False | 99.671% |
| HGCA ovary | 7,137 | 1,154,611 | 51,859 | False | 99.716% |
| HGCA skin | 26,476 | 1,154,611 | 59,422 | False | 99.665% |
| HGCA stomach | 18,710 | 1,154,611 | 33,091 | False | 99.780% |
| HGCA uterus | 8,475 | 1,154,611 | 72,572 | False | 99.611% |
